# CSF1R-dependent macrophages in the salivary gland are essential for epithelial regeneration following radiation-induced injury

**DOI:** 10.1101/2022.06.12.495803

**Authors:** John G. McKendrick, Gareth-Rhys Jones, Sonia S. Elder, Ella Mercer, Marlene S. Magalhaes, Cecilia Rocchi, Lizi M. Hegarty, Amanda L. Johnson, Christoph Schneider, Burkhard Becher, Clare Pridans, Neil Mabbott, Zhaoyuan Liu, Florent Ginhoux, Marc Bajenoff, Rebecca Gentek, Calum C. Bain, Elaine Emmerson

**Affiliations:** The Centre for Regenerative Medicine, The University of Edinburgh, 5 Little France Drive, Edinburgh, EH16 4UU, UK; The Centre for Inflammation Research, The University of Edinburgh, Queen’s Medical Research Institute, 47 Little France Drive, Edinburgh, EH16 4TJ, UK; The Centre for Reproductive Health, The University of Edinburgh, Queen’s Medical Research Institute, 47 Little France Drive, Edinburgh, EH16 4TJ, UK; Institute of Physiology, University of Zurich, Zurich, Switzerland; Institute of Experimental Immunology, University of Zurich, Zurich, Switzerland; The Roslin Institute & Royal (Dick) School of Veterinary Studies, The University of Edinburgh, Easter Bush Campus, Midlothian, EH25 9RG, UK; Shanghai Institute of Immunology, Department of Immunology and Microbiology, Shanghai Jiao Tong University School of Medicine, Shanghai 200025, China; Singapore Immunology Network, Agency for Science, Technology and Research, Singapore 138648, Singapore.; Translational Immunology Institute, SingHealth Duke-NUS Academic Medical Centre, Singapore, Singapore.; Centre d’Immunologie de Marseille-Luminy, Aix Marseille Université UM2, INSERM, U1104, CNRS UMR7280, 13288 Marseille, France

## Abstract

The salivary glands often become damaged in individuals receiving radiotherapy for head and neck cancer, resulting in xerostomia, or chronic dry mouth. This leads to detrimental effects on their health and quality of life, for which there is no regenerative therapy. Macrophages are the predominant cell type in the salivary glands and are attractive therapeutic targets due to their unrivalled capacity to drive tissue repair and regeneration. Yet, the nature and role of macrophages in salivary gland homeostasis and whether or not they contribute to tissue repair/regeneration following injury is not well understood. Here, we have used single cell RNA-seq, multi-parameter flow cytometry and fluorescence microscopy to map the heterogeneity of the salivary gland macrophage compartment throughout development and following radiation-induced injury. We show that there are highly dynamic changes in the composition of the salivary gland macrophage compartment with age, in part due to changes in the ontogeny of these cells, determined using a suite of complementary fate mapping systems. A combination of mutant mice and antibody blockade demonstrates that salivary gland macrophages are dependent on CSF1, but not IL-34 or GM-CSF, for their development and maintenance. Finally, using an *in vivo* model of radiation-induced salivary gland injury combined with a novel *Mafb*-specific depletion system, we demonstrate an essential role for macrophages. Without macrophages the clearance of cells with DNA damage, and effective tissue repair following such injury, is severely comprised. Our data, therefore, indicate a strong case for exploring the therapeutic potential of manipulating macrophages in order to promote tissue repair and thus minimise salivary gland dysfunction after radiotherapy.

## Introduction

Therapeutic radiation remains a mainstay for cancer treatment. While recent advances in the delivery of radiation aim to minimise off-target side-effects, healthy tissues that lie within the therapeutic field often also receive high doses of radiation, leading to cellular damage and organ dysfunction [1]. Thus, the salivary glands are often inadvertently damaged following radiotherapy for head and neck cancer, and this results in xerostomia, or chronic dry mouth [2]. While considerable efforts have been made to understand the side effects of radiation injury on the salivary glands, there is presently no regenerative therapy for this debilitating condition [1].

Macrophages have long been considered key cells in the tissue repair process and, in recent years, the idea of macrophages as therapeutic targets following radiation injury has become a realistic possibility [3]. However, macrophages are incredibly plastic cells that can adopt phenotypic and functional states depending on signals received from their environment, and the nature of these signals is known to change across the course of injury and repair [reviewed in 4, 5]. Thus, understanding the composition of tissue macrophage compartments, and whether this changes following injury, is key to determining whether macrophages could be targeted therapeutically. Application of single cell technologies, such as single cell RNA sequencing (scRNAseq), have revealed the true diversity of macrophages across and within tissues in both mouse and human [6–8]. Diversity can arise from discrete environmental imprinting in distinct anatomical locales, but also from changes in the ontogeny of macrophages. For instance, it is now clear that many tissue macrophages arise from embryonic progenitors that seed tissues during development [reviewed in 9]. However, the capacity of these embryonic-derived macrophages to persist in adulthood appears to be niche-specific. For example, while brain microglia derive exclusively from embryonic progenitors, macrophages in the choroid plexus are replaced by haematopoietic stem cells (HSC)-derived monocytes [10]; and although all gut macrophages are initially derived from embryonic sources, those in the mucosa are replaced by HSC-derived monocytes while muscularis macrophages appear to be relatively long-lived [11–14]. Importantly, macrophages of differing origins have been shown to play functionally-distinct roles in the context of infection and fibrosis in the lung [14, 15], in cardiac regeneration after myocardial infarction [16], and in the CNS in response to systemic endotoxin [17].

While the diversity and ontogeny of macrophages across most tissues has been described [6], those in the salivary glands remain relatively poorly defined [18]. To address this, we used immunophenotyping, confocal microscopy and scRNAseq to show that the mouse submandibular salivary gland (SMG) contains at least two populations of macrophages defined by their expression of CD206, CD163 and MHCII. Using a combination of lineage tracing approaches, we show that while there is a contribution of embryonic precursors to the SMG macrophage compartment during development, these embryo-derived macrophages are displaced in the late embryonic and neonatal stages by HSC-derived macrophages, which require low-rate replenishment by monocytes throughout life. We show that all SMG macrophages rely on signalling via the colony stimulating factor 1 receptor (CSF1R) for their development and maintenance. We show that radiation-induced injury leads to alterations in the composition of the macrophage pool and that although immediate post-radiation replenishment occurs through in-situ self-renewal, radiation treatment accelerates long-term macrophage replenishment from monocytes. Lastly, we use a *Mafb*-based depletion technique to demonstrate an indispensable role for macrophages in the repair and functionality of the SMG after irradiation. Together, these data highlight the integral role of macrophages during salivary gland repair.

## Results

### CSF1R-dependent macrophages dominate the SMG immune compartment

First, we set out to characterise the macrophage compartment of the naïve murine submandibular gland (SMG), the most well studied of the three major salivary glands, using a combination of flow cytometric analysis and multicoloured immunofluorescence. Amongst CD45^+^ cells, we could identify a large population of F4/80 expressing cells that made up the majority of leukocytes (**Fig. 1A**). Further phenotyping showed that these F4/80^+^ cells expressed low levels of CD11b and CD45, but high and uniform expression of CD172a (SIRPa) in the adult SMG (**Fig. 1A, B**). These F4/80^+^ cells also expressed high levels of CD64, a marker considered to be expressed by tissue macrophages [19], but lacked expression of Flt3, which is routinely used to define conventional dendritic cells (cDC) [20] (**Fig. 1B**). Immunofluorescent analysis showed that F4/80^+^ macrophages exist throughout the murine SMG epithelia, and surround both acini, identified by their distinctive circular structure and confirmed by staining with the acinar-specific marker aquaporin-5 (AQP5), and ducts, identified by their closely packed nuclei in a tubular arrangement and confirmed by strong expression of E-cadherin (ECAD) (**Fig. 1C**). SMG macrophages also expressed high levels of CSF1R, determined by using a novel CSF1R reporter mouse in which a cassette containing FusionRed fluorescent protein is inserted into the *Csf1r* locus [21]. To determine if SMG macrophages were reliant on CSF1R signalling for their development and/or maintenance, we assessed mice deficient in a super-enhancer region of the CSF1 receptor, termed the *fms*-intronic regulatory element (FIRE) (*Csf1r*^ΔFIREΔ/FIRE^ mice), which selectively impacts CSF1R expression and tissue macrophage development in a variety of tissues [22]. Adult *Csf1r*^ΔFIRE/ΔFIRE^ mice had significantly fewer F4/80^+^ SMG macrophages compared with *Csf1r*^+/+^ and *Csf1r*^+/ΔFIRE^ littermates (**Fig. 1D, E**). In contrast, *Csf2ra*^−/−^ mice, which are deficient for the alpha subunit of the receptor for granulocyte-macrophage colony stimulating factor (GM-CSF; also known as CSF2), had normal density of SMG macrophages compared with *Csf2ra*^+/+^ littermates (**Fig. S1A**), ruling out a role for GM-CSF in the development/maintenance of these cells. To complement our findings in *Csf1r* ^ΔFIREΔ/FIRE^ mice, we administered an anti-CSF1R blocking antibody (AFS98) to *Cx3cr1*^+/gfp^ mice for three days before assessing macrophage abundance. Treatment caused a marked reduction of F4/80^+^CX3CR1^+^ SMG macrophages, not seen in recipients of the isotype control antibody (**Fig. 1F**). There are two known ligands for the CSF1R: CSF1 and IL-34. While *Il34*-deficient mice lack Langerhan’s cells and microglia [23, 24], analysis of adult *Il34*^LacZ/LacZ^ mice showed no difference in the abundance of SMG macrophages when compared with *Il34*^LacZ/+^ littermates (**Fig. 1G**). Therefore, and in keeping with previously published studies [25, 26], our results demonstrate conclusively that SMG macrophages depend on the CSF1-CSF1R axis for both their development and maintenance.

**Figure 1:**
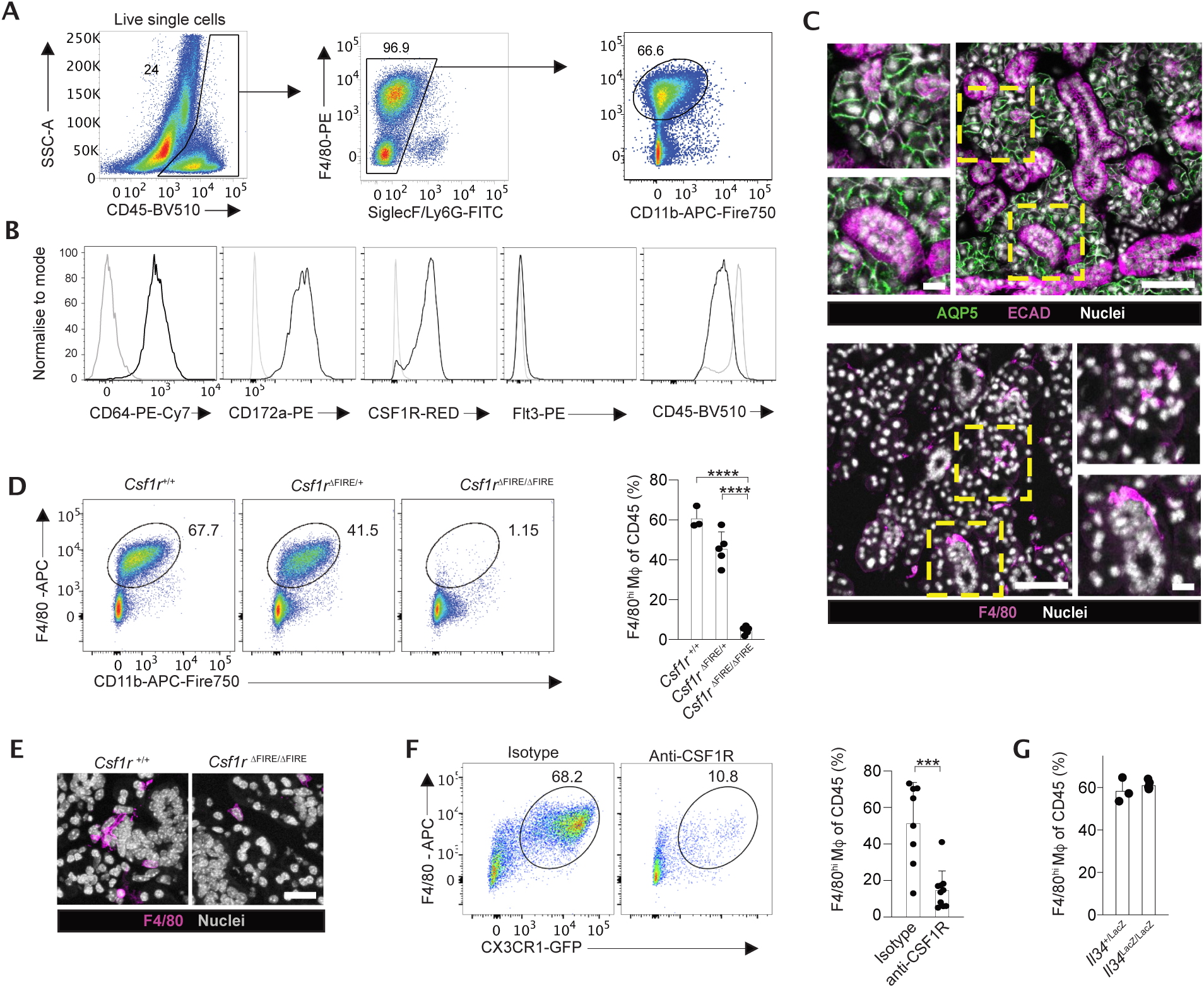
CSF1R-dependent macrophages dominate the SMG immune compartment. **A.** Gating strategy for the identification of F4/80^+^ macrophages in the unmanipulated submandibular salivary gland (SMG). **B.** Representative expression of CD64, CD172a, CD45 and Flt3 by F4/80^+^ macrophages obtained from SMG from unmanipulated adult C57BL/6J mice and expression of CSF1R-Red by F4/80^+^ macrophages obtained from SMG from unmanipulated adult *Csf1r*^FRed^ mice. Data are from one of two independent experiments. Grey histograms represent fluorescence minus one (FMO) controls or background fluorescence of SMG macrophages from *Csf1r*^WT^ mice in the case of CSF1R-Red expression. **C.** Representative expression of aquaporin 5 (AQP5) and E-cadherin (ECAD) (upper panels) and F4/80 (lower panels) in SMG tissue from unmanipulated adult C57BL/6J mice. Scale bars in large panel = 100μm, magnified insets = 10μm. **D.** Representative expression of F4/80 and CD11b by CD45^+^Lin^−^ live leukocytes from the SMG of unmanipulated adult *Csf1r*^ΔFIRE/ΔFIRE^ mice and their *Csf1r*^+/+^ and *Csf1r*^ΔFIRE/+^ littermates. Graphs show the frequency of macrophages of CD45^+^ leukocytes. Data are from 3 (*Csf1r*^+/+^), 5 (*Csf1r*^ΔFIRE/+^) or 5 (*Csf1r*^ΔFIRE/ΔFIRE^) mice per group and are pooled from 2 independent experiments. ***p<0.001 (one-way ANOVA with post-hoc Tukey *Q* test) and error bars represent the SD. **E.** Representative expression of F4/80 in SMG tissue of unmanipulated adult *Csf1r*^ΔFIRE/ΔFIRE^ mice and their *Csf1r*^+/+^ littermates. Scale bar = 10μm. **F.** Representative frequency of F4/80^+^CD11b^lo^ macrophages in SMG of adult *Cx3cr1*^+/gfp^ mice administered anti-CSF1R (AFS98) or isotype control for 3 days. Data are from 8 mice per group and are pooled from two independent experiments. ***p<0.001 (unpaired Student’s *t* test) and error bars represent the SD. **G.** Frequency of F4/80^+^CD11b^lo^ macrophages in SMG of adult *Il34*^LacZ/LacZ^ mice compared with *Il34*^+/LacZ^ littermate controls. Data are from 3 (*Il34*^+/LacZ^) or 5 (*Il34*^LacZ/LacZ^) mice per group and are from 1 representative experiment of 2.

### Developmentally-related heterogeneity in macrophages exists in the SMG

To allow further characterisation of these cells in an unbiased manner, we next performed single cell RNA-seq (scRNA-seq) of non-granulocytic, non-lymphocytic F4/80^+^ myeloid cells (CD3^−^CD19^−^ NK1.1^−^Ly6G^−^SiglecF^−^) obtained from unmanipulated adult SMG using the 10X Chromium platform (**Fig. S2A**). Because we also sequenced endothelia and epithelia in this analysis, we identified macrophages, and their putative subsets, on the basis of *Adgre1* (encoding F4/80) and *Csf1r* (**Fig. S2B)**, and re-clustered these cells (**Fig. 2A**). This revealed four clusters of cells. All subsets expressed *Cx3cr1* and *C1qa*, as well as the macrophage-specific transcription factor *Mafb* (**Fig. 2B, c**). Cluster 3 was defined by expression of genes associated with the cell cycle, including *Mki67, Tubb5* and *Top2a*, suggesting these cells represent proliferating macrophages (**Fig. 2B, C**), a finding consistent with low G2/M and S phase gene expression (**Fig. S2C**) [27]. Cluster 0 was defined by higher expression of *Cd81*, *Trem2*, *Apoe*, *Cd63 and Hexb,* whereas Cluster 1 expressed higher levels of *Cd14, Cd83, Ccl3, Ccl4, Cxcl2* and *Cxcl10* (**Fig. 2B, C**). Cluster 2 appeared to be very distinct when compared with the other clusters, and was defined by expression of *Folr2, Mrc1* (encoding CD206; also known as mannose receptor) and *Cd163* (**Fig. 2A-C**). To validate this heterogeneity, we assessed expression of folate receptor β (FRβ, encoded by *Folr2*), CD206, CD163, CD14, CD63 and MHCII by flow cytometry. We found a small but distinct population of CD206^+^ macrophages that co-expressed FRβ and CD163 in the homeostatic adult SMG (**Fig. 2D**) and demonstrated the presence of CD206^+^ and CD163^+^ macrophages in the SMG by immunofluorescence (**Fig. 2E** and **Fig. S2D**). Moreover, CD206^+^ macrophages lacked expression of CD11c, which was expressed by CD206^−^ macrophages in the adult SMG (**Fig. 2D**). While we detected expression of surface CD14 and MHCII, and intracellular CD63 by flow cytometry, these did not identify discrete subsets of cells amongst CD206^−^ macrophages (**Fig. 2D**), suggesting that these clusters shared a surface phenotype but exhibit clear transcriptional differences. Finally, by comparing with other tissue macrophages including those from brain, colon, lung, spleen and liver [28], we were able to define the SMG macrophage transcriptional ‘signature’, and show that despite exhibiting features associated with microglia, including low expression of CD45 and high expression of *Hexb*, SMG macrophages have most similarity with colonic macrophages (**Fig. S2E-G**).

**Figure 2:**
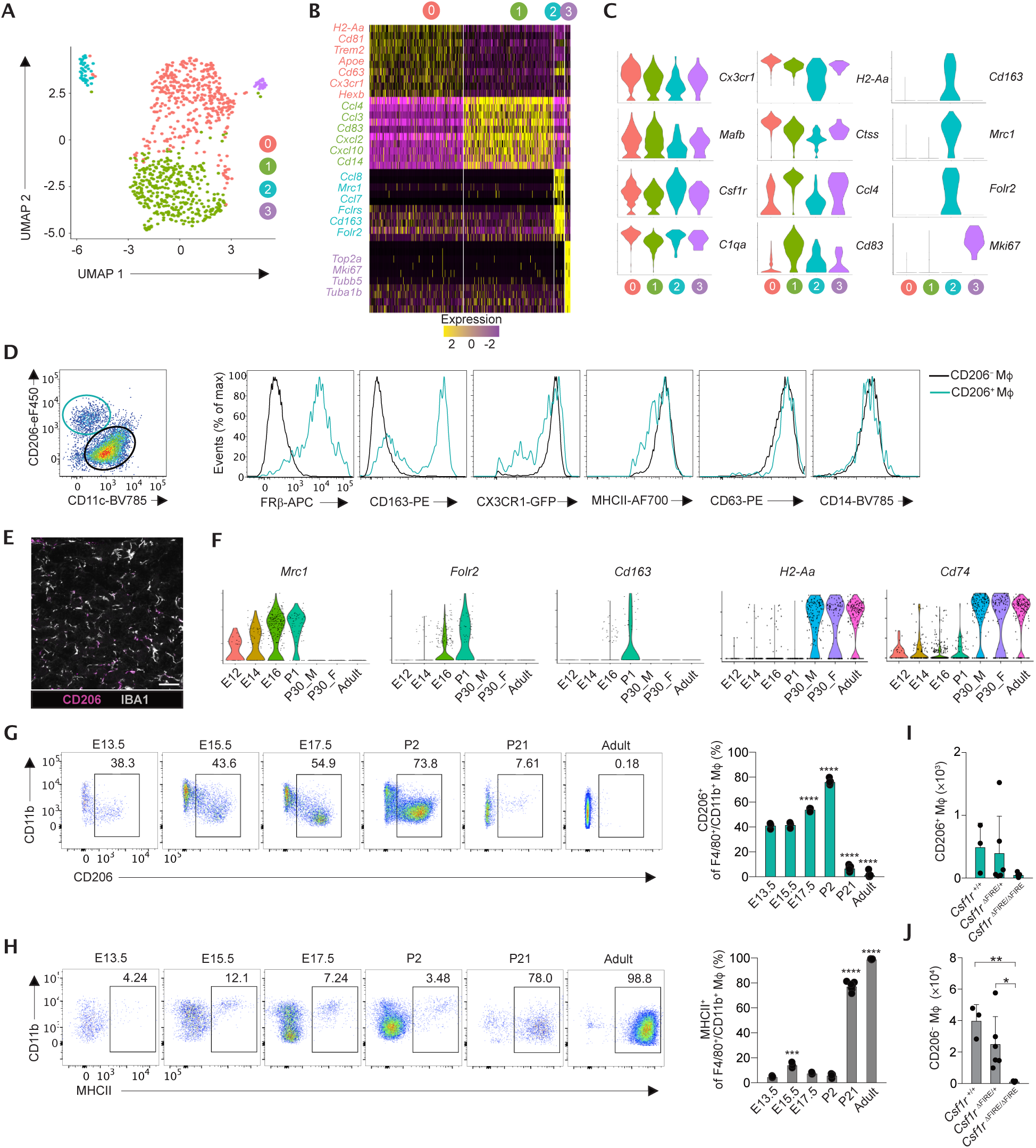
Developmentally-related heterogeneity in macrophages exists in the SMG. **A.** UMAP dimensionality reduction analysis of *Adgre1*^+^*Csf1r*^+^ cells from scRNA-seq of SMG of unmanipulated adult male C57BL/6J mice. **B.** Heatmap showing the top 20 most differentially expressed genes by each cluster defined in **A.** and annotated to show genes of particular interest. **C.** Violin plots showing gene expression of curated pan-macrophage and cluster defining genes from **B**. **D.** Representative expression of CD206, CD11c, FRβ, CD163, MHCII, CD63, CD14 by F4/80^+^ macrophages obtained from SMG from unmanipulated adult C57BL/6J mice. Data are from one of two independent experiments. **E.** Representative expression of CD206 and IBA1 in SMG tissue of unmanipulated adult C57BL/6 mice. Scale bar = 50μm. **F.** Feature plots showing expression of *Mrc1*, *Folr2, Cd163, H2-Aa* and *Cd74* by *Adgre1*^+^*Csf1r*^+^ cells across a developmental time course from [30]. **G-H.** Representative expression of CD206 (**G**) or MHCII (**H**) by F4/80^+^ macrophages from SMG of mice at the indicated ages. Graphs show the mean frequency of CD206^+^ macrophages of total F4/80^+^ macrophages. Data are from 3 (E13.5, 15.5, 17.5, P2), 4 (P21) and 3 (adult) mice per group and are pooled from 2 independent experiments. ****p<0.0001 (one-way ANOVA with post-hoc Tukey *Q* test) compared to E13.5 and error bars represent the SD. **I-J.** Absolute number of CD206^+^ (**I**) and CD206^−^ (**J**) macrophages in the unmanipulated adult SMG of *Csf1r*^τιFIRE/τιFIRE^ mice and their *Csf1r*^+/+^ and *Csf1r*^τιFIRE/+^ littermates. Data are from 3 (*Csf1r*^+/+^), 5 (*Csf1r*^τιFIRE/+^) or 5 (*Csf1r*^τιFIRE/τιFIRE^) mice per group and are pooled from 2 independent experiments. *p<0.05, **p<0.01 (one-way ANOVA with post-hoc Tukey *Q* test) and error bars represent the SD.

To determine if the heterogeneity seen in the adult SMG, and in particular the presence of *Mrc1^+^Folr2^+^* macrophages, was apparent throughout SMG development, we examined macrophage heterogeneity in embryonic and neonatal SMG. Gland ontogenesis is initiated at embryonic day 11.5 (E11.5), epithelial branching begins at E13, and luminisation and terminal differentiation occurs at E16, forming an organ capable of secretion by postnatal day 7 (P7) [reviewed in 29]. We examined the profile of SMG macrophages across the developmental time course using a publicly available scRNA-seq ‘atlas’ dataset of whole SMG [30]. Using a similar approach to the above, we extracted macrophages from this dataset on the basis of *Adgre1* and *Csf1r* (**Fig. S2H, I**). While there were relatively low numbers of macrophages in this dataset compared with our scRNA-seq analysis, we could detect differences between macrophages from E14, P1 and adult SMG. Macrophages from E14 and P1 clustered more closely together than adult SMG, suggesting they are more alike (**Fig. S2I**). Strikingly, expression of *Mrc1* and *Folr2* appeared to increase during the embryonic stages until P1, yet expression was low or almost absent by 30 days of age and in mature adults (**Fig. 2F**). Conversely, *H2-Ab* and *Cd74* (the invariant chain of the MHCII complex), which appeared to define Clusters 0 and 1 in our scRNA-seq dataset, were expressed at negligible levels during development, but expressed at high levels by macrophages in the adult SMG (**Fig. 2F**). Confocal microscopy confirmed the presence of CD163^+^ and CD206^+^ macrophages during embryonic development (**Fig. S2J, K)**. Again, we used flow cytometry to validate these scRNA-seq data, showing that CD206^+^ macrophages were present in the embryonic SMG, and came to dominate the neonatal SMG, whilst only forming a small minority of SMG macrophages by adulthood (**Fig. 2G**). In parallel, expression of MHCII appeared to be induced in the postnatal period, with ∼75% of macrophages expressing MHCII by 3 weeks of age and is essentially present in all macrophages by adulthood (**Fig. 2H**), consistent with postnatal acquisition of MHCII by macrophages in other tissues [6]. Importantly, analysis of adult *Csf1r*^ϕλFIRE/ϕλFIRE^ mice showed that bothCD206-defined populations of SMG macrophages rely on the CSF1R for their development/maintenance (**Fig. 2I-J**). Thus, dynamic changes to the composition of the CSF1R-dependent SMG macrophage compartment occur in the late embryonic, neonatal and juvenile periods.

### Postnatal switch in the ontogeny of SMG macrophages

Given the presence of macrophages in the embryonic SMG and the dynamic changes seen during the neonatal and juvenile periods, we next assessed the ontogeny of SMG macrophages using multiple, complementary fate mapping approaches that together delineate their progenitor sources and homeostatic turnover from the adult bone marrow. We first used *Cdh5*^CreERT2/+^.*Rosa26*^CAG-LSL-tdT/+^*.Cx3cr1*^GFP/+^ mice to fate map cells during embryonic development. Yolk sac-derived progenitors and definitive haematopoietic stem cells (HSCs) are produced at different sites and developmental stages [31], but due to their endothelial origin, both can be labelled in the *Cdh5*^Cre-ERT2^ model [32] via administration of 4-hydroxytamoxifen at either E7.5 (yolk sac) or E10.5 (HSC) (**Fig. 3A**). As expected, brain microglia were highly labelled in offspring of mothers pulsed at E7.5 (**Fig. 3B, C**). In contrast, blood monocytes from the same mice showed only very low levels of labelling. Importantly, we found that, similar to blood monocytes, the total SMG macrophage compartment showed minimal contribution of cells labelled at E7.5 when analysed in newborn mice (P1) or in adult mice (**Fig. 3B, C**). However, analysis of CD206-defined subsets revealed differences in labelled efficiency. Consistent with the presence of CD206^+^ in the embryonic gland, labelling was markedly higher in this fraction compared with CD206^−^ macrophages. By adulthood there was no longer a difference in the presence of yolk-sac derived macrophages in the gland, suggesting that these cells are largely displaced by HSC-derived cells. Analysis of the offspring of mothers pulsed at E10.5 supported this finding. In these mice microglia were very poorly labelled, but circulating monocytes were efficiently labelled, consistent with their derivation from HSCs (**Fig. 3D**). SMG macrophages were found to have equivalent labelling to that seen in monocytes. We did find a small but significant difference in labelling between CD206-defined macrophage subsets, with lower labelling in CD206^+^ macrophages (**Fig. 3D**), suggesting that the rate of replenishment may differ between these subsets.

**Figure 3:**
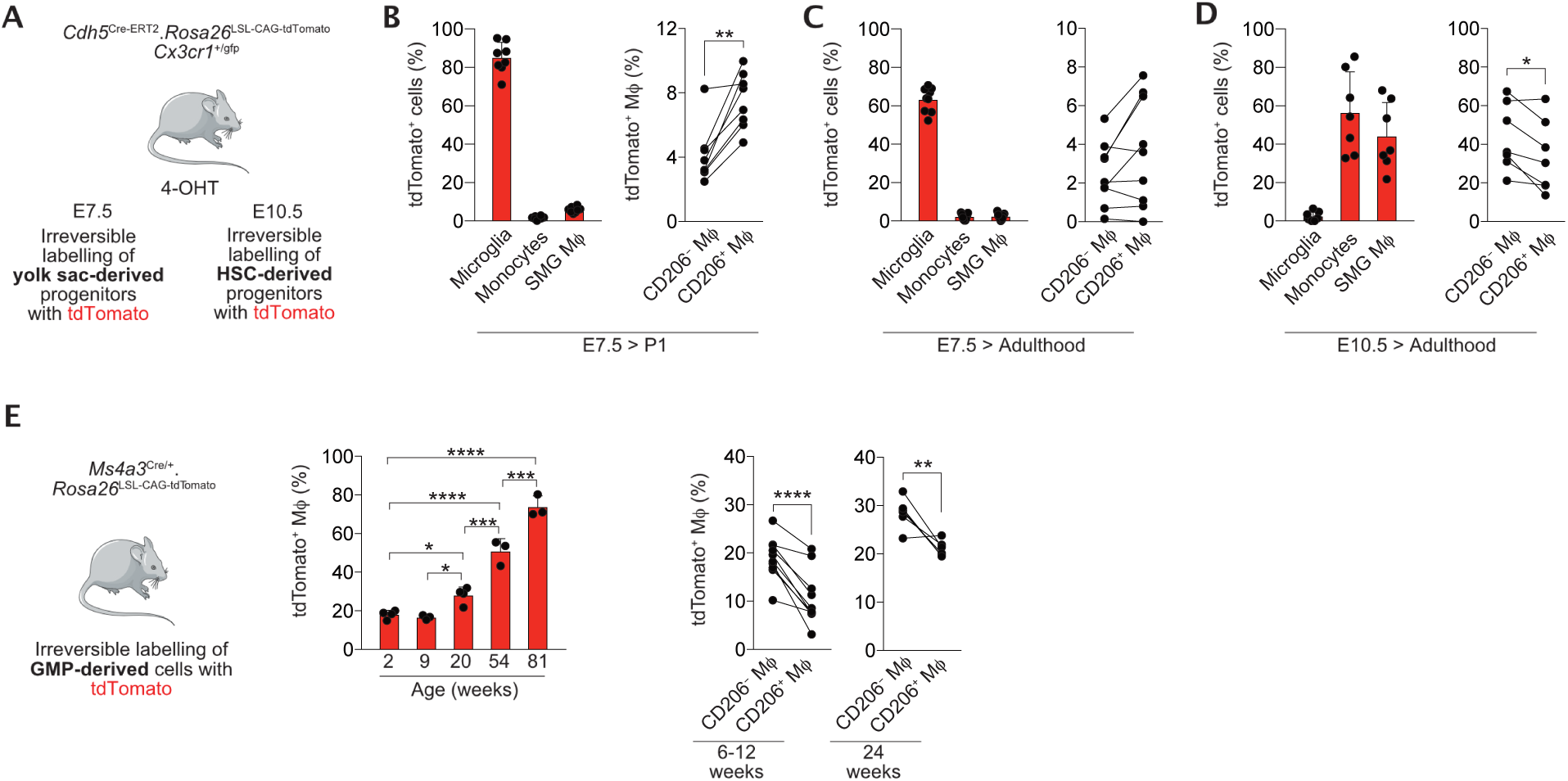
Postnatal switch in the ontogeny of SMG macrophages. **A.** Experimental scheme for fate mapping with *Cdh5*^Cre-ERT2^.*Rosa26*^LSL-CAG-tdTomato^*.Cx3cr1*^+/gfp^ mice. **B-C.** The frequency of tdTomato^+^ cells amongst brain microglia, blood monocytes and SMG F4/80^+^CD11b^lo^ macrophages as a total population (*left*) or as CD206-defined subsets (*right*) in the neonatal (P1) (**B**) or adult (**C**) offspring of mice pulsed with 4-hydroxytamoxifen (4-OHT) at embryonic day 7.5 (E7.5). Data are from 3 independent litters (**B**) or 8 adult mice pooled from 2 experiments (**C**). **p<0.01 (paired Student’s *t* test). **D.** The frequency of tdTomato^+^ cells amongst brain microglia, blood monocytes and SMG F4/80^+^CD11b^lo^ macrophages as a total population (*left*) or as CD206-defined subsets (*right*) in adult offspring of mice pulsed with 4-OHT at embryonic day 10.5 (E10.5). Data are from 7 mice pooled from 2 experiments. *p<0.05 (paired Student’s *t* test). **E.** Description of *Ms4a3*^Cre/+^.*Rosa26*^LSL-CAG-tdTomato^ mice. Graph shows the frequency of tdTomato^+^ cells amongst total F4/80^+^CD11b^lo^ macrophages (*left*) or CD206-defined subsets (*right*) obtained from *Ms4a3*^Cre/+^.*Rosa26*^LSL-CAG-tdTomato^ mice at the indicated ages. Data are from n=4 mice for 2- and 20 week time points, and n=3 for all other timepoints. **p<0.01 (unpaired Student’s *t* test). In all graphs symbols represent individual mice and error bars represent the standard deviation (SD).

To test this directly, we next assessed the contribution of monocytes to SMG at different life stages by examining *Ms4a3*^Cre/+^.*Rosa26*^CAG-LSL-tdTomato^ mice in which granulocyte-monocyte progenitors (GMPs) and their progeny are labelled irreversibly with tdTomato fluorescent protein (**Fig. 3E**). Consistent with their derivation from BM GMPs, circulating monocytes are labelled with high efficiency (>95%) in adult *Ms4a3*^Cre/+^.*Rosa26*^LSL-CAG-tdTomato/+^ mice [33]. In contrast, brain microglia show negligible labelling in the same mice [33]. Longitudinal analysis of these mice showed that there was slow, but progressive accumulation of tdTomato label in the SMG compartment (**Fig. 3E**), consistent with the accumulation of monocyte-derived cells. In keeping with the differences seen between CD206-defined subsets in the *Cdh5*-based fate mapping, we found the CD206^+^ macrophages were labelled to a lower extent in *Ms4a3*^Cre/+^.*Rosa26*^LSL-CAG-tdTomato/+^ mice than their CD206^−^ counterparts in young adults (**Fig. 3E**).

Taken together, these data demonstrate that embryo-derived macrophages initially seed the SMG, but are entirely replaced by HSC-derived cells, most likely in the neonatal period. These cells become long-lived macrophages, although they are replenished at a low rate by BM-derived monocytes.

### Irradiation injury alters the composition and longevity of SMG macrophages

Having established the heterogeneity and replenishment kinetics of SMG macrophages during homeostasis, we next used our well-validated mouse model of irradiation injury, where the neck and SMGs are irradiated with a single dose of 10Gy while the rest of the body is shielded (**Fig. 4A**), to characterise how these cells respond following irradiation. As previously reported in the murine sublingual gland [34], transcriptional analysis of whole SMG tissue confirmed early cellular apoptosis and epithelial damage, with a significant increase in pro-apoptotic *Bax* expression at days 1 and 3 post-irradiation, which had returned to baseline levels by day 7 (**Fig. 4B**). In addition, expression of *Aqp5,* which encodes the water channel protein AQP5, was essentially absent at day 3, before recovering by day 7 and day 28, with a peak of expression at day 14. Consistent with exposure to radiation, overall proliferation, as measured by *Ki67* expression at tissue level, was reduced at day 1 but elevated at day 3 before returning to baseline thereafter. In parallel, we found a trend towards lower levels of *Csf1r* mRNA in SMG following radiation, which was restored to previous levels by day 14 (**Fig. 4B**), suggesting radiation may alter the abundance and/or the composition of the macrophage pool. Consistent with this, enumeration of macrophages by confocal microscopy showed that radiation treatment led to a reduction in the absolute numbers of SMG macrophages, which recovered by day 7 (**Fig. 4C**). Following radiation injury, F4/80^+^ macrophages were found to preferentially localise to cells displaying signs of DNA damage, marked by expression of 53BP1 [35] (**Fig. 4C**). Interestingly, flow cytometric analysis of SMG showed that radiation injury caused little or no changes in expression of canonical markers, including F4/80, MHCII and CD11b (**Fig. 4D, E**). In parallel, there was no measurable recruitment of neutrophils or monocytes, suggesting that radiation treatment did not elicit a marked inflammatory response (**Fig. S3A**). In support of this, expression of mRNA for the pro-inflammatory cytokines IL-1β, IL-6 and TNF did not significantly change across the irradiation time course (**Fig. S3B**). To assess if the transcriptional fingerprint of macrophages changed in the context of irradiation, we collected SMG at days 0, 3 and 28 post-irradiation, and undertook scRNA-seq analysis (**Fig. 4F-I**). We chose these timepoints to span peak injury (day 3) and injury resolution (day 28). As before, we artificially selected specific cells for analysis by sorting macrophages, epithelial cells and endothelial cells. Macrophages were again identified on the basis of *Adgre1* (F4/80) and *Csf1r* expression, and re-clustered (**Fig. S3C**). Again, this revealed 4 clusters (termed A, B, C, D). Clusters A and B were the two largest clusters and aligned with Cluster 0 and 1, respectively, in our earlier analysis (**Fig. 4F**). Cluster C represented the CD163^+^CD206^+^ population referred to as Cluster 3 in the previous analysis, while Cluster D denotes a population of cycling macrophages, defined by expression of *mKi67, Stmn1* and *Top2a* (**Fig. 4F**) and aligned with Cluster 2 in the previous analysis. The most striking effect of radiation treatment was the almost complete loss of Cluster D at day 3 post-radiation, consistent with the known effects of radiation interrupting cell proliferation. Notably, by day 28 the relative abundance of Cluster D had recovered to levels seen at steady state (**Fig. 4F, G**) and we confirmed this pattern using flow cytometry by measuring Ki67 expression (**Fig. 4H** and **Fig. S3D**). Pairwise gene expression analysis revealed that certain changes in gene expression were present across clusters, for example upregulation of *Cxcl2, Atf3* and *Nfkbia* at day 3 (**Fig. S3E**). However, cluster-specific differences were apparent. For instance, Cluster A, which expresses higher levels of *Tnf* and *Il1b* during physiological conditions, showed upregulation of these transcripts following radiation treatment (**Fig. 4I**). Another prominent feature of this cluster following radiation was expression of *Cdkn1a* (**Fig. 4I**), which encodes cyclin-dependent kinase 1A (also known as p21), a negative regulator of cell cycle. Expression of CDKN1A is known to confer radio-resistance to epidermal Langerhans cells [36] and Kupffer cells [37], and this could explain the relative preservation of this subset. Cluster A also showed elevated expression of *Nlrp3* (**Fig. 4I**), which is part of the inflammasome and detects products of damaged cells including ATP and uric acid [38], suggesting these cells may play a particular role in the recognition and elimination of damaged cells. Following radiation Cluster B displayed elevated expression of *Lgals3* (Galectin-3) and *Tyrobp* (DAP12) (**Fig. 4I**), genes that are involved in macrophage activation and recruitment [39–41]. Post-radiation Cluster C showed heightened expression of the monocyte chemoattractants *Ccl8, Ccl7, Ccl2 and Ccl6* (**Fig. 4I**). In line with previously published data, this population likely acts to recruit other immune cells, while the release of CC chemokines may also confer radioresistance and preservation of this subset [42].

**Figure 4:**
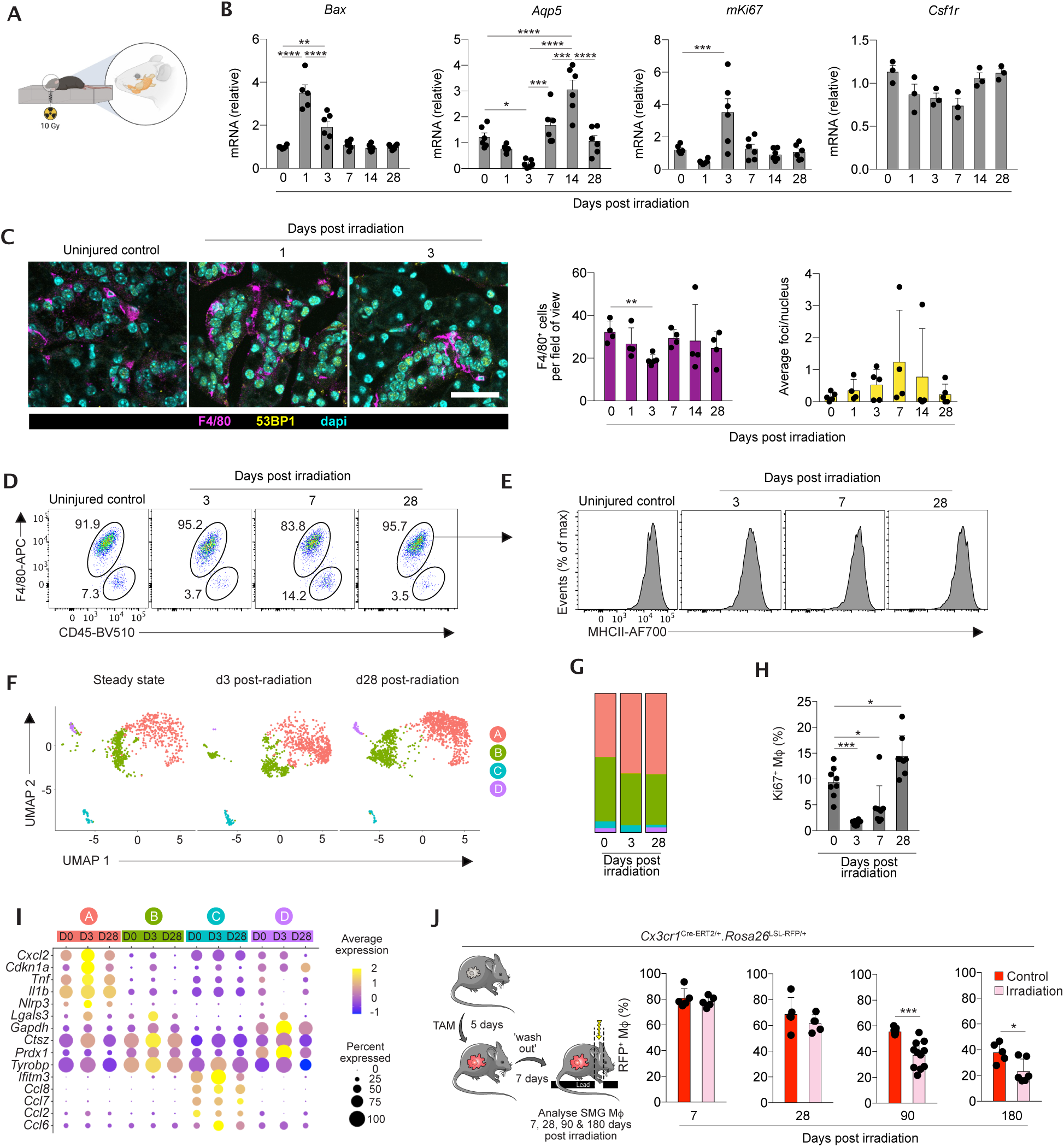
Irradiation injury alters the composition and longevity of SMG macrophages. **A.** Schematic of targeted radiation **B.** qPCR analysis of *Bax*, *Aqp5*, *Ki67* and *Csf1r* mRNA in total SMG tissue at the indicated time points following radiation induced injury. Data are normalised to mRNA levels in unmanipulated naïve (d0) SMG tissue. Data are from 3-6 mice pooled from 2 independent experiments. *p<0.05, **p<0.01, ***p<0.001, ****p<0.0001 (One-way ANOVA followed by Tukey *Q* post-hoc test). **C.** Representative expression of F4/80 and 53BP1 in the SMG of uninjured mice or mice irradiated 1 or 3 days earlier. Scale bar = 25μm. Graphs show the enumeration of F4/80^+^ and 53BP1^+^ cells at the indicated time points following radiation. Data obtained from three fields of view from non-sequential sections from 4-5 mice per timepoint. **p<0.01 (One-way ANOVA with Tukey *Q* post-hoc testing). **D.** Representative expression of F4/80 and CD45 obtained from SMG of uninjured mice or mice irradiated 3, 7 or 28 days earlier. **E.** Representative expression of MHCII by F4/80^+^CD11b^lo^ macrophages obtained from SMG of uninjured mice or mice irradiated 3, 7 or 28 days earlier. **F.** UMAP dimensionality reduction analysis of scRNA-Seq data from 3,478 *Adgre1*^+^ and *Csf1r*^+^ macrophages split by timepoint. **G.** Proportion by total number of each macrophage sub-clusters defined in **F** from SMG of unmanipulated mice or mice irradiated 3 or 28 days earlier. **H.** Frequency of Ki67^+^ F4/80^+^CD11b^lo^ macrophages in the SMG at 3, 7 and 28 days following radiation exposure compared with untreated mice (d0). Data are from 8 mice per time point pooled from two independent experiments. *p<0.05, **p<0.01 ***p<0.001 (One-way ANOVA followed by Tukey *Q* post-hoc testing). **I.** Bubble plot showing expression of selected genes by macrophage clusters from unmanipulated SMG (d0) or SMG at day 3 (d3) or day 28 (d28) following radiation treatment. **J.** Experimental scheme for fate mapping CX3CR1^+^ cells in adult *Cx3cr1*^Cre-ERT2/+^.*Rosa26*^LSL-RFP^ mice. Graphs shows the frequency of RFP^+^ cells amongst F4/80^+^CD11b^lo^ SMG macrophages obtained from *Cx3cr1*^Cre-ERT2/+^.*Rosa26*^LSL-RFP^ mice administered tamoxifen by oral gavage for 5 days and analysed at the indicated time points following targeted radiation. Data are 5-11 mice pooled from 3 independent experiments. *p<0.05, ***p<0.001 (unpaired Student’s t test). In all graphs symbols represent individual mice and error bars represent the SD.

The changes in the macrophage compartment seen through scRNAseq analysis and, in particular, the almost complete loss of proliferating macrophages at the acute time point analysed (day 3) prompted us to evaluate whether radiation altered macrophage replenishment kinetics. To this end, we performed pulse-chase fate mapping using *Cx3cr1*^Cre-ERT2/+^.*Rosa26*^LSL-RFP/+^ mice by administering tamoxifen for five days, followed by a ‘wash out’ period of one week to ensure circulating classical monocytes were no longer labelled (**Fig. 4J**). Mice were then subjected to targeted irradiation or left untreated, and loss of RFP signal amongst macrophages measured as a rate of macrophage replenishment from RFP^−^ monocytes. Interestingly, while radiation treatment led to an initial reduction in the absolute number of macrophages (**Fig. 4C**), which recovered within the first week, such treatment had no effect on the frequency of RFP-expressing macrophages at this stage. This suggests that the replenishment of macrophages at this point appears to occur independently of blood monocytes (**Fig. 4J**). When examined at day 28, the repair phase, although loss of RFP^+^ macrophages could be detected compared with day 7, this loss was similar between the groups (**Fig. 4J**). However, by 3 months there was significantly greater loss of RFP-labelled SMG macrophages in irradiated mice compared with controls, a phenotype that remained evident at 6 months post-irradiation (**Fig. 4J**).

Taken together these data suggest that there are subset-specific responses to radiation treatment, that initial recovery in macrophage numbers occurs through in situ proliferation and that elevated macrophage replenishment from the blood monocytes occurs in the long term.

### SMG macrophages are essential for epithelial regeneration following irradiation injury

Finally, to demonstrate a role for SMG macrophages in tissue repair after irradiation injury, we used an *in vivo* depletion system. In lieu of systems which selectively target SMG macrophage subsets (CD206^+^CD11c^−^ and CD206^−^CD11c^lo-hi^), we used *Mafb*^Cre/+^:*Cx3cr1*^LSL-DTR/+^ mice, in which all *Cx3cr1*-expressing macrophages are rendered susceptible to diphtheria toxin (DTx), in order to temporally and selectively deplete macrophages in the SMG (**Fig. 5A**). As shown earlier, all subsets of SMG macrophages express both *Mafb* and *Cx3cr1* (**Fig. 2C**). We first demonstrated that we could achieve efficient depletion of SMG macrophages using this model. Following two doses of DTx, we observed a significant reduction in F4/80^+^ macrophages in the otherwise healthy SMG (**Fig. 5B**). Using this methodology, we then combined macrophage depletion with irradiation injury. Immunofluorescence analysis revealed fewer Ki67^+^ cells in the absence of macrophages at 3 days post-radiation (**Fig. 5C**), while CASP3^+^ apoptotic cells were elevated (**Fig. 5C**). In parallel, where macrophage depletion had been efficient, as defined by areas devoid of F4/80^+^ cells, DNA damage marked by 53BP1^+^ foci in the cell nuclei was elevated at day 7 post-radiation (**Fig. 5D**), consistent with the role of macrophages in clearing damaged and dying cells.

**Figure 5:**
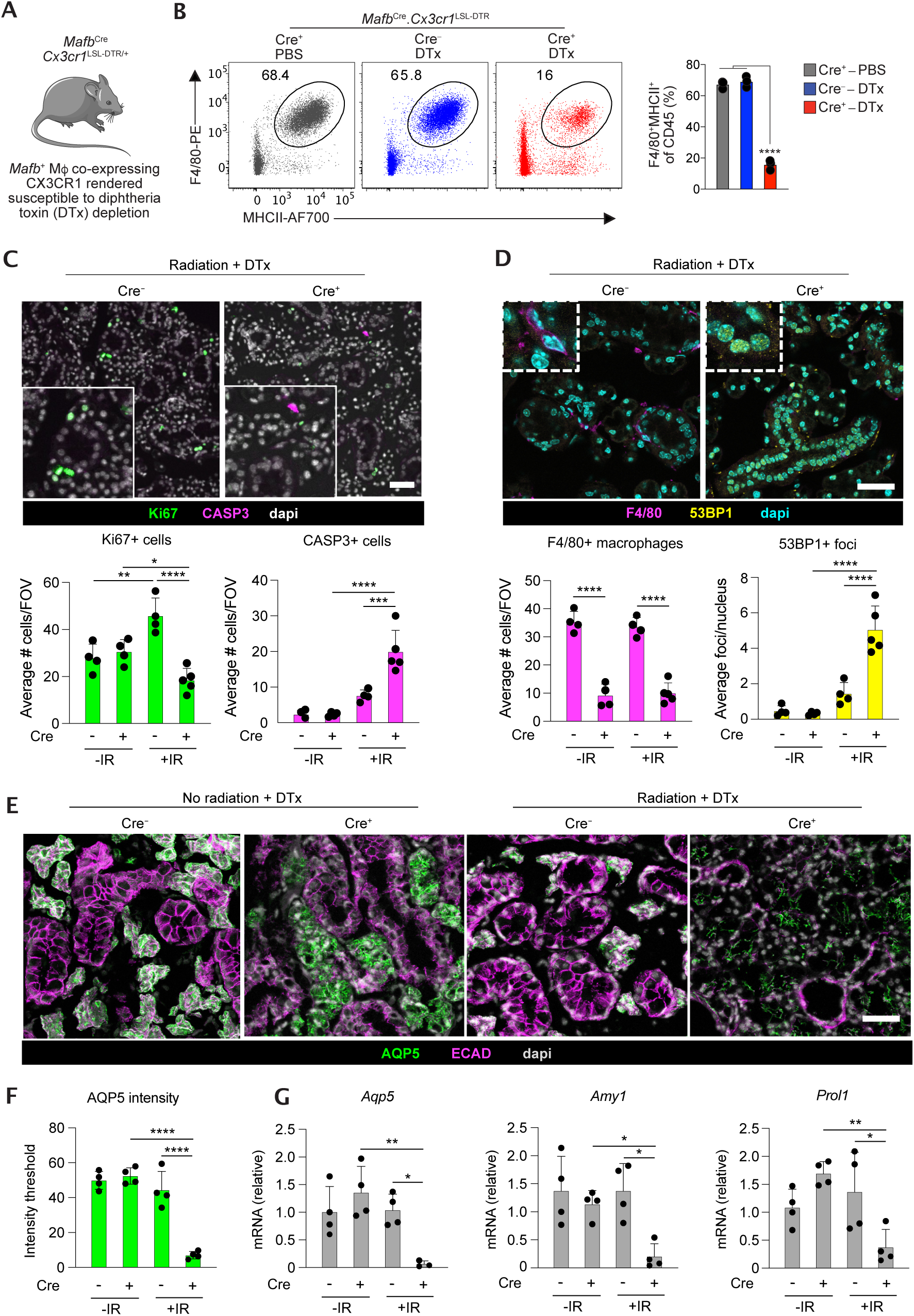
SMG macrophages are essential for epithelial regeneration following irradiation injury. **A.** Schematic of the *Mafb*^Cre/+^.*Cx3cr1*^LSL-DTR/+^ mouse model. **B.** Representative expression of F4/80 and MHCII by CD45^+^SiglecF^−^Ly6G^−^ leukocytes obtained from *Mafb*^Cre/+^.*Cx3cr1*^LSL-DTR/+^ mice or *Mafb*^+/+^.*Cx3cr1*^LSL-DTR/+^ littermates administered diphtheria toxin (DTx) or saline (PBS). Data are from 3 mice from one of two independent experiments performed. ****p<0.0001 (One-way ANOVA followed by Tukey *Q* post-hoc testing). Symbols represent individual mice and error bars represent the SD. **C.** Representative immunofluorescent images of SMG stained for Ki67 and CASP3 in *Mafb*^Cre/+^.*Cx3cr1*^LSL-DTR/+^ mice or *Mafb*^+/+^.*Cx3cr1*^LSL-DTR/+^ littermates administered diphtheria toxin (DTx) or saline for 5 days before being exposed to 0Gy or 10Gy irradiation, and analysed 3 days after irradiation. Scale bar = 25μm. Graphs show the enumeration of Ki67^+^ and CASP3^+^ cells. Data obtained from three fields of view from non-sequential sections from 4 mice per group. *p<0.05, **p<0.01, ***p<0.001, ****p<0.0001 (One-way ANOVA with Tukey *Q* post-hoc testing). **D.** Representative immunofluorescent images of SMG stained for F4/80 and 53BP1 in *Mafb*^Cre/+^.*Cx3cr1*^LSL-DTR/+^ mice or *Mafb*^+/+^.*Cx3cr1*^LSL-DTR/+^ littermates administered diphtheria toxin (DTx) or saline for 5 days before being exposed to 0Gy or 10Gy irradiation, and administered diphtheria toxin (DTx) or saline for a further 3 days before being analysed 7 days after irradiation. Scale bar = 25μm. Graphs show the enumeration of Ki67^+^ and CASP3^+^ cells. Data obtained from three fields of view from non-sequential sections from 4 mice per group. ****p<0.0001 (One-way ANOVA with Tukey *Q* post-hoc testing). **E.** Representative immunofluorescent images of SMG stained for AQP5 and ECAD in *Mafb*^Cre/+^.*Cx3cr1*^LSL-DTR/+^ mice or *Mafb*^+/+^.*Cx3cr1*^LSL-DTR/+^ littermates exposed to 0Gy or 10Gy irradiation and administered diphtheria toxin (DTx) or saline from day 17 onwards, every 2 days and analysed 28 days after irradiation. Scale bar = 25μm. **F.** Enumeration of AQP5 intensity in *Mafb*^Cre/+^.*Cx3cr1*^LSL-DTR/+^ mice or *Mafb*^+/+^.*Cx3cr1*^LSL-DTR/+^ littermates exposed to 0Gy or 10Gy irradiation and administered diphtheria toxin (DTx) or saline from day 17 onwards, every 2 days and analysed 28 days after irradiation. Data obtained from three fields of view from non-sequential sections from 4 mice per group. ****p<0.0001 (One-way ANOVA with Tukey *Q* post-hoc testing). **G.** qPCR analysis of *Aqp5*, *Amy1* and *Prol1* (Mucin-10) mRNA in total SMG tissue in *Mafb*^Cre/+^.*Cx3cr1*^LSL-DTR/+^ mice or *Mafb*^+/+^.*Cx3cr1*^LSL-DTR/+^ littermates exposed to 0Gy or 10Gy irradiation and administered diphtheria toxin (DTx) or saline from day 17 onwards, every 2 days and analysed 28 days after irradiation. Data are normalised to mRNA levels in SMG tissue *Mafb*^+/+^.*Cx3cr1*^LSL-DTR/+^ littermates exposed to 0Gy. Data are from 4 mice per group. *p<0.05, **p<0.01 (One-way ANOVA followed by Tukey *Q* post-hoc test).

Studies in mice have shown that after irradiation injury the salivary gland tissue goes through an initial regenerative phase, which ultimately fails in the long term [43–45], mirroring what occurs in individuals following radiotherapy [46–48]. We therefore extended our studies to examine the roles of macrophages in the initial regenerative phase by determining the effects of macrophage depletion at the point at which many indices of damage and inflammation have subsided, and initial repair is under way (day 28). To this end, we administered DTx every other day from day 17 to day 27 and analysed the SMG at day 28 using the parameters described above. This revealed that macrophage depletion hindered the restoration of salivary gland structure following irradiation injury. Specifically, while the structural and functional markers AQP5 (acinar cells) and ECAD (all epithelia, but high in ductal cells) have returned to a level comparable with uninjured SMG by 28 days in the presence of macrophages (**Fig. 5E**), macrophage depletion (Cre^+^ +DTx) led to aberrant patterning of AQP5 and irregular ductal structures 28 days after injury (**Fig. 5E** and **Fig. S4A**). Given that AQP5 is necessary for water transfer and is the only aquaporin to play a major role in the salivary secretion process [49], the lack of AQP5 here demonstrates that macrophage depletion severely impacts on secretory function. Indeed, ablation alone without irradiation injury also results in structural alterations (**Fig. 5E**), albeit to a lesser extent, indicating that SMG macrophages may also support homeostatic maintenance of SG structural cells. While macrophage depletion did not have an effect on overall gland weight (**Fig. S4B**), expression of genes which represent function, including aquaporin-5 (*Aqp5*), Amylase 1 (*Amy1*) and Mucin-10 (*Prol1*) and the epithelial adhesion marker E-cadherin (*Cdh1*) were significantly reduced at day 28, while the acinar specification marker *Sox10* was unchanged (**Fig. 5G** and **Fig. S4C**).

The above results demonstrate clearly that macrophages are crucial at both early and later stages of repair of the SMG following irradiation injury.

## Discussion

Macrophage heterogeneity and ontogeny in many tissues has been studied extensively in recent years, yet the salivary glands have remained largely unexplored despite containing a dense network of macrophages [50]. Here we have used a combination of approaches to demonstrate that there is considerable heterogeneity within the salivary gland macrophage compartment throughout the life course, resulting in part from changes in the ontogeny of these cells. Furthermore, we show that salivary gland macrophages are crucial for effective tissue repair following radiation-induced injury.

Previous work has utilised scRNA-seq to chart the cellular composition of the salivary gland [30, 51]. However, most of these studies have sought to create a cellular ‘atlas’ of the salivary gland and thus have lacked sufficient resolution to comprehensively and specifically profile the macrophage compartment. We therefore performed targeted scRNA-seq of the macrophage compartment, along with epithelial and endothelial cells, although we did not analyse these non-haematopoietic cells in this study. This provided sufficient resolution to identify four clusters of macrophages which fell into three phenotypically discrete subsets identifiable by flow cytometry: CD206^+^, CD206^−^ and proliferating macrophages (Ki67^+^). CD206-defined macrophages differed in their expression of a range of surface markers, including FRβ, CD163 and CD11c, as well as expression of chemokines, supporting the idea that these are distinct macrophage subsets. Clusters 0 and 1 appeared to represent discrete transcriptional states of the CD206^−^CD11c^lo-hi^ subset. Our characterisation is consistent with work by the Stein group where CD11c-YFP reporter mice were used to visualise SMG macrophages, and where most but not all SMG macrophages expressed YFP in the adult SMG [50]. The hypothesis that SMG macrophages can be partitioned on the basis of CD11c expression is also consistent with a recent study [52], although we propose that positive identification of CD11c^−^ macrophages on the basis of CD206 (or FRβ) expression is a superior way to characterise the SMG compartment. Our transcriptional profiling also supports the notion that CD11c^+^ macrophages interact with T cells in the SMG. In particular, the constitutive expression of *Cxcl10* by CD206^−^CD11c^+^, but not CD206^+^CD11c^−^, macrophages may support CXCR3-dependent clustering of tissue resident memory CD8^+^ T cells adjacent to macrophages in the SMG [50].

Longitudinal analysis across the life course demonstrated that CD206^+^ macrophages dominate the developing and neonatal SMG but form only a minor fraction of the adult SMG macrophage compartment, a finding that we validated through analysis of a publicly available scRNA-seq dataset. Again, this is consistent with the study by Lu *et al.* [52] where CD11c^+^ macrophages were shown to accumulate in the postnatal SMG. Previous work has failed to reach a consensus on the replenishment kinetics of SMG macrophages. For instance, while global *Ccr2* deficiency has been used to support the idea that SMG macrophages require no replenishment from classical CCR2-dependent monocytes [53], others have used CCR2 antagonism to reach opposite conclusions [52]. Our study is the first to use state-of-the-art genetic lineage tracing models to document the ontogeny of SMG macrophages across the lifecourse in both healthy tissue and in the context of injury. This combination of models demonstrated that although embryonic progenitors contribute to the initial seeding of the SMG macrophage compartment, these are displaced by HSC-derived macrophages, and low-level contribution of monocytes is needed to maintain the SMG macrophage compartment during adulthood, a finding reproduced using *Ms4a3^Cre^* reporter mice from three institutions. In the spectrum of macrophage replenishment [54], SMG macrophages appear to have similar kinetics to red pulp splenic macrophages [33]. Notably, the rate of replenishment of CD206^+^ macrophages appeared to be lower than the CD206^−^CD11c^+^ subset. This could reflect a difference in growth factor dependence, although our analysis of *Csf1r*^ΔFIRE/ΔFIRE^ mice suggests that both CD206-defined subsets depend on CSF1R for their development and maintenance. We also ruled out a role for GM-CSF and the CSF1R ligand IL-34 in SMG macrophage homeostasis. Taken together with previous work showing a deficit of salivary gland macrophages in *Csf1r* knockout rats [26], similar to that in the *Csf1*^op/op^ mouse [25], which has a naturally occurring inactivating mutation in the *Csf1* gene, our data demonstrate a key role for the CSF1-CSF1R axis in controlling SMG macrophage homeostasis. This is particularly pertinent given that: CSF1 promotes branching morphogenesis of the mouse SMG [55]; CSF1 is expressed in response to Hedgehog (Hh) signalling in the SMG [56]; and that acinar cells are reported to express CSF1 [57]. Together these indicate epithelial-macrophage communication. Our developmental studies suggest that there is a shift in the niche during the embryonic and neonatal period, as macrophages migrate into the surrounding mesenchyme and appear in closer proximity to the epithelia. Given that acinar cell maturation occurs during the post-natal period, their provision of CSF1 may result in some of the postnatal changes in SMG macrophages. Moreover, this macrophage-epithelial communication is likely to be bidirectional as depletion of SMG macrophages in *Mafb*^Cre/+^.*Cx3cr1*^LSL-DTR/+^ mice in the absence of radiation treatment led to alterations in the epithelial compartment. Understanding the nature of this crosstalk will be a key aim of future work.

In this study we show, for the first time, that radiation injury leads to a shift in SMG macrophage subsets and a striking loss of a proliferative subset. Surprisingly, macrophage repopulation following injury occurred independently of monocytes and was associated with elevated levels of proliferation by residual macrophages. Whether all macrophages possess identical capacity to proliferate, or if proliferative subsets may exhibit “stemlike” characteristics [58] is still not known and will require novel transgenic systems. Since we observed a loss of these macrophages in the early stages after radiation injury, and given the demonstration that proliferative resident macrophages are essential for islet cell proliferation in the pancreas [59], understanding more about these subsets will be integral for future studies. It is clear that in the long-term macrophage replenishment is accelerated by radiation treatment, suggesting that radiation may cause long-term changes to the macrophage niche, making it less able to support macrophage longevity.

Our model of radiation-induced injury allowed us to assess macrophage function in the immediate response (day 3) and during tissue repair (day 28). Crucially, we demonstrate that depletion of resident CX3CR1^+^ macrophages in the initial days after irradiation injury leads to an accumulation of DNA damage, failure of epithelial recovery and a loss of tissue functionality. Complementary experiments showed that macrophages play a key role in supporting tissue repair at day 28 after injury. In particular, our data indicate a role for macrophages in orchestrating epithelial repair and restoration of the secretory function of acinar cells. Our fate mapping experiments support the premise that these temporal functions are performed by the same cells; however, whether distinct subsets play differential roles in the response to radiation-induced injury remains unclear, and will require the generation of novel, subset-specific targeting strategies.

Here we have performed comprehensive characterisation of the SMG macrophage compartment in health, across development, and in the context of radiation-induced damage. We have demonstrated that discrete macrophage subsets exist in the healthy SMG and that there are both acute and long-term effects of radiation on the transcriptional fingerprint of SMG macrophages and their replenishment kinetics. Understanding the factors that drive macrophage specialisation, how these change following radiation, and how the niche is altered, will all be vital in order to therapeutically target macrophage response to irradiation damage.

## Materials and Methods

### Mouse studies

All procedures were approved by the UK Home Office and performed under PPLs PB5FC9BD2, PP0860257, PP1871024. Mice of both sexes were used. Sex and age of mice is noted for each experiment in the relevant figure legend. Transgenic mice used in this study are listed in Table 1.

**Table 1.**
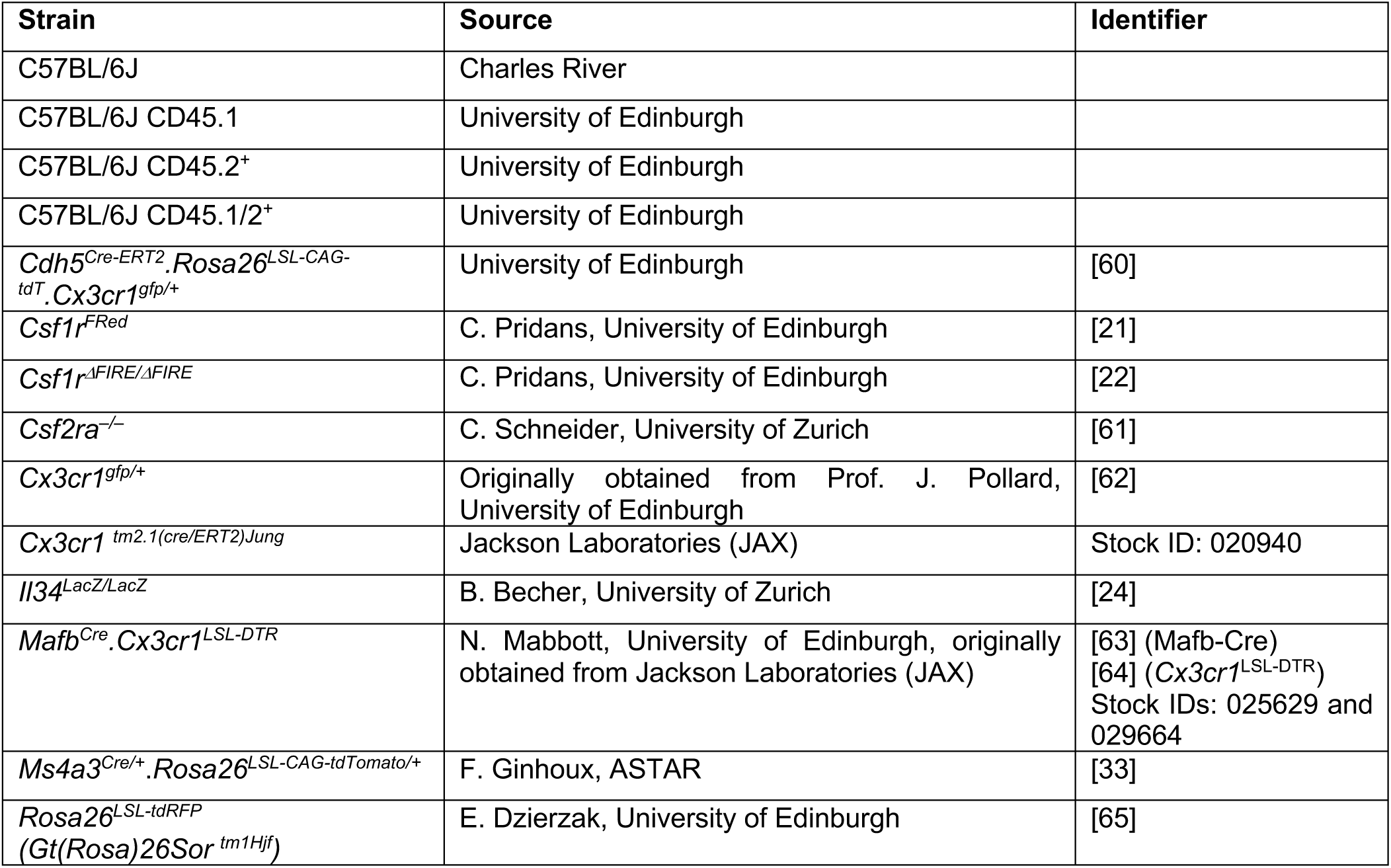
Mouse strains used in this study, their source and relevant identifiers.

### Lineage tracing studies

Labelling of CX3CR1^+^ cells was achieved by administering *Cx3cr1*^Cre-ERT/+^.*Rosa26*^LSL-RFP/+^ mice with 5mg tamoxifen (Sigma Aldrich) in 100μl sunflower oil (Sigma Aldrich) by oral gavage on 5 consecutive days, followed by 7 days of no treatment (‘washout’) before further manipulation (e.g. irradiation). For *Cdh5^Cre-ERT2^*fate mapping, WT female mice aged 6-10 weeks were subjected to timed matings with *Cdh5^Cre-ERT2+/-^* or *Cdh5^Cre-ERT2+/+^ Rosa^tdT/tdT^ Cx3cr1^gfp/gfp^* males. Successful mating was judged by the presence of vaginal plugs the morning after, which was considered 0.5 days post-conception. For induction of reporter recombination in the offspring, a single dose of 4-hydroxytamoxifen (4OHT; 1.2mg) was delivered by intraperitoneal (i.p) injection to pregnant females at E7.5 or E10.5. To counteract the adverse effect of 4OHT on pregnancy, 4OHT was supplemented with progesterone (0.6mg). In cases when females could not give birth naturally, pups were delivered by C-section and cross-fostered with CD1 females.

### Anti-CSF1R experiments

Anti-CSF1R antibody (AFS98) was generated as previously described [66, 67]. Male C57BL/6 mice were administered with intravenous (i.v) injection of 2mg/mL anti-CSF1R antibody over three consecutive days (250 µg per mouse per day). Mice were assessed the day after the final administration.

### Genetic depletion of Cx3cr1^+^ cells

Conditional depletion of *Cx3cr1^+^ cells* was achieved by injecting *Mafb^Cre/+^.Cx3cr1^LSL-DTR/+^* mice with 200ng Diphtheria toxin (DTx; Sigma Aldrich, D0564) in 100 μl sterile saline intraperitoneally (IP) 3x per week. For homeostatic assessment of depletion, mice were examined 24hrs after the final DTx dose. When combined with radiation treatment, mice received DTx every other day for 5 days before irradiation (single dose of 10Gy) and were examined at days 3 and 7 post-irradiation, or mice underwent irradiation before receiving DTx every other day for 10 days from day 17 post-irradiation, and were examined at day 28 post-irradiation.

### Radiation-induced SG injury

Mice were anaesthetised using 1mg/kg Medetomidine Hydrochloride (Dormitor) and 75mg/kg ketamine (Ketavet) in 0.9% saline (Thermo Fisher Scientific). Mice were irradiated using a single ^137^Cs source in a Shepherd Mark-I-68A ^137^Cs irradiator (JL Shepherd & Associates), with only the neck exposed and the rest of the head and body shielded with lead, as previously described [34]. Non-irradiated control mice were not anaesthetised. After a 20 minute period of anaesthesia, mice were given 1mg/kg of reversal agent Antisedan and were allowed to recover on a heat pad before returning to normal housing. Subsequently, mice were provided with soft diet and DietGel® (Clear H_2_O) *ad libitum*. Mice were euthanised at 1, 3, 7, 14 or 28 days or 3 or 6 months post-irradiation.

### Developmental studies

Embryonic SMGs were collected from embryos at E12.5, E13.5, E14.5, E15.5, E17.5 and postnatal animals at P2, P7 and P21, following timed matings between male and female C57BL/6 mice. Successful mating was confirmed by the presence of a vaginal plug and the morning of discovery deemed E0.5. SMG explants were dissected as previously described [68] and used for immunofluorescent analysis or flow cytometry.

### Cell dissociation Embryonic SMG

SMGs were pooled from multiple mice for E12.5, E13.5, E14.5 (n=3 per sample) given their small size, in order to obtain sufficient material. E15.5, E17.5, P2, P7 and P21 glands were analysed individually. SMGs were digested in 500μl RPMI-1640 containing 5% fetal calf serum (FCS; Sigma Aldrich), 6µl Collagenase-II (23mg/mL) (Sigma Aldrich), 6µl hyaluronidase (40mg/mL) (Sigma

Aldrich) and 37µl 0.1M CaCl_2_ for 10 minutes in a shaking incubator at 100 rpm at 37°C. Tissue was subsequently centrifuged at 400G for 5 minutes at 4°C. The supernatant was discarded and the pellet resuspended in 500μl of HBSS (Lonza) containing 1% BSA (Sigma Aldrich) and 2mM EDTA (Sigma Aldrich) and then passed through a 20µm filter capped 5mL FACs tube using a 5mL syringe with a 25G needle.

### Adult SMG

Each SMG pair (approx. 160mg) was mechanically minced in 2mL of RPMI-1640 containing 5% fetal calf serum (FCS; Sigma Aldrich), 25µl Collagenase-II (23mg/mL) (Sigma Aldrich), 25µl hyaluronidase (40mg/mL) (Sigma Aldrich) and 125µl 0.1M CaCl_2_ using a GentleMACS machine, program A.01 in a GentleMACS C-Tube. Tissue was incubated in a shaking incubator at 100 rpm at 37°C for 60 minutes and subsequently centrifuged at 400G for 5 minutes at 4°C. The supernatant was discarded and the pellet was resuspended in 2mL of RPMI-1640 containing 5% FCS. The solution was filtered through a 70µm nylon mesh (ThermoFisher) and centrifuged at 400G for 5 minutes at 4°C. The cell pellet was resuspended in 1mL of 1x red blood cell lysis buffer (Abcam) for 5 minutes on ice before centrifugation at 400G for 5 minutes at 4°C. The cell pellet was resuspended in 1mL of pre-warmed (37°C) trypsin +0.25% EDTA and incubated at 37°C for 5 minutes before trituration. This step was repeated 3 times until single cells were evident. The solution was centrifuged at 400G for 5 minutes at 4°C and the cell pellet was resuspended in 1mL of Hank’s Balanced Salt Solution (HBSS; Lonza) containing 1% BSA (Sigma Aldrich) and 2mM EDTA (Sigma Aldrich) and then passed through a 20µm filter capped 5mL FACS tube using a 5mL syringe with a 25G needle.

### Isolation of brain microglia

For analysis of microglia, single cell suspensions were obtained from brain tissues via a combination of enzymatic digest and mechanical dissociation. Brain tissue was finely minced with scissors and digested in RPMI medium (Invitrogen) containing 2% FCS (Sigma Aldrich), 0.1 mg/ml DNase I (Sigma Aldrich), 200U/ml Collagenase I (GIBCO), 1mg/ml Dispase II (Roche) and 0.1mg/ml Liberase DL (Roche). Samples were incubated at 37°C for 30-45 minutes under agitation (900 rpm), with regular trituration. Microglia were identified as CD45^dim^ CD64^+^ CD11b^+^ Cx3cr1^+^.

### Flow cytometry and fluorescence-activated cell sorting (FACS)

Equal numbers of cells were stained with 1:1000 anti-CD16/32 (2.4G2; Biolegend) in 100µl of HBSS containing 1% BSA and 2mM EDTA (termed FACS buffer hereafter) for 15 minutes, to reduce non-specific antibody binding to receptors for IgG. Cells were subsequently stained with conjugated antibodies (**Table 2**) for 30 minutes at 4°C in the dark. Samples were washed with FACS buffer and centrifuged at 400G for 5 minutes at 4°C before resuspension in 300µl of FACS buffer. Single stain controls were prepared using OneComp Beads (ThermoFisher). Fluorescence minus one (FMO)

**Table 2.** Antibodies used for flow cytometry.

| Antibody | Clone | Supplier | Cat # | Dilution | RRID |
| --- | --- | --- | --- | --- | --- |
| Rat CD11b APC/Fire 750 | M1/70 | Biologend | 101262 | 1:200 | AB_2572122 |
| Rat CD11b PE | M1/70 | Biologend | 101207 | 1:200 | AB_312790 |
| Armenian Hamster CD11c BV785 | N418 | Biologend | 117336 | 1:200 | AB_2565268 |
| Rat CD14 Superbright 600 | Sa2-8 | Invitrogen | 63-0141-82 | 1:200 | AB_2762769 |
| CD14 BV785 | Sa14-2 | Biologend | 123337 | 1:200 | AB_2888880 |
| Rat CD16/CD32 TruStain FcX | S17011E | Biologend | 156603 | 1:1000 | AB_2783137 |
| Rat CD45 APC-Cy7 | 30-F11 | BD Biosciences | 557659 | 1:200 | AB_396774 |
| Rat CD45 BV510 | 30-F11 | Biologend | 103138 | 1:200 | AB_2563061 |
| Mouse CD45.1 AF488 | A20 | Biologend | 110718 | 1:200 | AB_492862 |
| Mouse CD45.1 BV510 | A20 | Biologend | 110741 | 1:200 | AB_2563378 |
| Mouse CD45.2 AF700 | 104 | Biologend | 109822 | 1:200 | AB_493731 |
| CD63 PE | NVG-2 | Biologend | 143903 | 1:200 | AB_11203532 |
| Mouse CD64 BV421 | X54-5/7.1 | Biologend | 139309 | 1:200 | AB_2562694 |
| Rat CD86 FITC | GL1 | Invitrogen | 11-0862-85 | 1:200 | AB_465149 |
| Rat CD163 PE | S15049F | Biologend | 156704 | 1:200 | AB_2860724 |
| Mouse CD206 PE/Cy7 | MR6F3 | Invitrogen | 25-2061-82 | 1:200 | AB_2637424 |
| Rat CD206 eFluor 450 | MR6F3 | Invitrogen | 48-2061-80 | 1:200 | AB_2762720 |
| Rat CD209 PE | LWC06 | Invitrogen | 12-2092-82 | 1:200 | AB_657702 |
| Mouse CX3CR1 PE | SA011F11 | Biologend | 149005 | 1:200 | AB_2564314 |
| Mouse CX3CR1 Biotin | SA011F11 | Biologend | 149018 | 1:200 | AB_2565701 |
| Mouse CX3CR1 APC | SA011F11 | Biologend | 149007 | 1:200 | AB_2564491 |
| Rat F4/80 PE | BM8 | Biologend | 123110 | 1:200 | AB_893486 |
| Rat F4/80 PE/Cy5 | BM8 | Biologend | 123111 | 1:200 | AB_893494 |
| Rat F4/80 APC | BM8 | Invitrogen | 17-4801-82 | 1:200 | AB_2784648 |
| FRβ APC | 10/FR2 | Biologend | 153306 | 1:200 | AB_2721313 |
| Ki67 FITC | REA183 | Miltenyi | 130-117-803 | 1:200 | AB_2733584 |
| Mouse Ly6C APC eFluor 780 | HK1.4 | Invitrogen | 47-5932-82 | 1:200 | AB_2573992 |
| Rat Ly6C PerCP/Cy5.5 | HK1.4 | Biologend | 128012 | 1:200 | AB_1659241 |
| Rat Ly6G Biotin | 1A8 | Biologend | 127604 | 1:200 | AB_1186108 |
| Rat Ly6G FITC | 1A8 | Biologend | 127606 | 1:200 | AB_1236494 |
| Rat MerTK PE | 2B10C42 | Biologend | 151506 | 1:200 | AB_2617037 |
| Rat MHC II AF700 | M5/114.15.2 | Biologend | 107622 | 1:200 | AB_493727 |
| Rat MHC II PE/Cy5 | M5/114.15.2 | Invitrogen | 15-5321-82 | 1:200 | AB_468800 |
| Mouse SiglecF FITC | REA798 | Miltenyi | 130-112-178 | 1:200 | n/a |
| Streptavidin BV650 | n/a | Biologend | 405231 | 1:10000 | n/a |
| Rat Tim4 PE/Cy7 | RMT4-54 | Biologend | 130010 | 1:200 | AB_2565719 |

controls were prepared using cells. Counting beads (ThermoFisher) were included to calculate absolute numbers. Dapi (Sigma Aldrich) or SYTOX Green (ThermoFisher) was used as a dead cell marker. Samples were analysed using an LSRII (BD Biosciences). Data was analysed using FlowJo (V9 or V10).

### Tissue processing for histology

SMGs were fixed for 6-8 hours in 4% paraformaldehyde (PFA; Thermo Scientific) at room temperature, with constant mixing, followed by 3 x washes with phosphate buffered saline (PBS; Merck). After fixation, SMGs were processed for the generation of frozen sections, by incubating in increasing concentrations of sucrose (10% and 30%) before embedding in OCT (Leica). 12 μm sections were cut using a cryostat (Leica) and stored at −20 °C.

### Immunofluorescent analysis

Whole-mount salivary gland and tissue section immunofluorescence analysis have been previously described [34, 69]. In brief, tissue was fixed with 4% PFA if not previously fixed, and permeabilised with ice cold acetone/methanol (1:1) for 1 min. Tissue was blocked for 2 hours at room temperature with 5% BSA (Sigma Aldrich), 5% Donkey Serum (Merck) in 0.01% PBS-Tween-20. Salivary glands were incubated with primary antibodies overnight at 4 °C. Antibodies are listed in **Table 3**. Antibodies were detected using donkey Cy2-, Cy3- or Cy5-conjugated secondary Fab fragment antibodies (Jackson Laboratories) and nuclei stained using Hoechst 33342 (1:1000, Sigma Aldrich), and mounted using Prolong Gold anti-fade mounting media. Fluorescence was analysed using a Leica SP8 confocal microscope and NIH ImageJ software.

**Table 3.** Primary antibodies used for immunofluorescent staining.

| Antibody | Clone | Species | Supplier | Cat # | Dilution | RRID |
| --- | --- | --- | --- | --- | --- | --- |
| E-Cadherin | ECCD2 | Rat | Life Technologies | 13-1900 | 1:300 | AB_2533005 |
| AQP5<sup>o</sup> |  | Rabbit | Millipore | AB3559 | 1:200 | AB_2141915 |
| CD163 | TNKUPJ | Rat | Thermo Fisher | 14-1631-82 | 1:800 | AB_2716934 |
| CD206-APC | C068C2 | Mouse | Biolegend | 141708 | 1:200 | AB_10896057 |
| CD45 |  | Goat | R&D Systems | AF114 | 1:200 | AB_442146 |
| F4/80 | A3-1 | Rat | Abcam | ab6640 | 1:200 | AB_1140040 |
| IBA1 |  | Rabbit | Antibodies Online | ABIN2857032 | 1:200 |  |
| 53BP1 |  | Rabbit | Novus | NB100-304 | 1:1200 | AB_1659863 |
| Ki67 | SolA15 | Rat | Invitrogen | 14-5698-82 | 1:200 | AB_10854564 |
| CASP3 | 5A1E | Rabbit | Cell Signalling | 9664L | 1:300 | AB_2070042 |

<sup>o</sup> This antibody has been discontinued

### Histological cell counts

For immunofluorescent analysis, cells positively stained for markers were counted using ImageJ. 3 random fields of view per sample were taken on a Leica SP8 microscope at 40x magnification (Nyquist). Images were run through an ImageJ cell counting macro either as single images or, if the file was a z-stack, the middle image of the stack was used. Using the macro, images were split into individual channels and the appropriate channel was extracted. Positive cells, such as macrophages, were thresholded and counted according to their size using the “analyse particles” command. As a quality control an output file was saved where the counted macrophages were highlighted in green and could be manually checked/confirmed. The macro is included in the Supplemental Methods. Fluorescence intensity was measured in Image J using the Threshold function.

### Quantitative PCR analysis (qPCR)

RNA was isolated from whole tissue using the RNAqueous Micro Kit (Life Technologies). Total RNA samples were DNase-treated (Life Technologies) prior to cDNA synthesis (First Strand Synthesis Kit, ThermoFisher). SYBRgreen qPCR was performed using 5ng cDNA and primers designed using Primer3 and Beacon Designer software or described on PrimerBank (http://pga.mgh.harvard.edu/primerbank/). Primer sequences are listed in **Table 4**. Melt-curves and primer efficiency were determined as previously described [70]. Gene expression was normalized to the housekeeping gene *Gapdh* and to the corresponding experimental control. Reactions were run in triplicate.

**Table 4.**
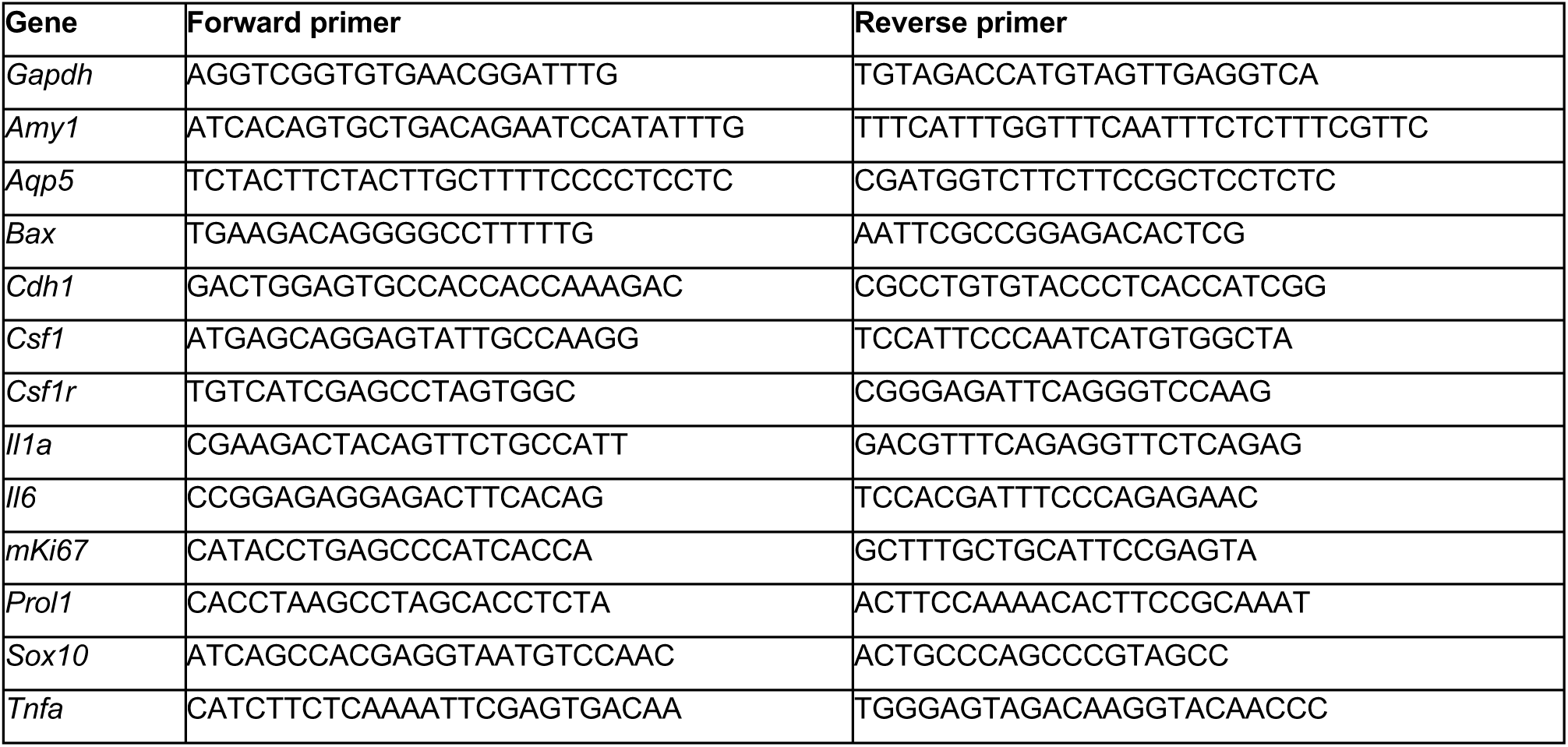
Primer sequences used for qPCR.

### Transcriptional profiling by scRNA-seq

Male C57BL/6 mice were irradiated (n=2 per group), as previously described and euthanised at 3 or 28 days post-irradiation. Non-irradiated control mice were not anaesthetised. Each SMG pair (approx. 160mg) was processed into a single cell suspension, as described for flow cytometry. Equal numbers of cells were stained with 1:1000 anti-CD16/32 (2.4G2; Biolegend) in FACS buffer for 15 minutes, to reduce non-specific antibody binding to receptors for IgG. Cells were subsequently stained with conjugated antibodies (**Table 5**) for 30 minutes at 4°C in the dark. Epithelial cells (EpCAM^+^), endothelial cells (CD31^+^) and macrophages (CD45^+^CD11b^+^F4/80^+^SiglecF^-^Ly6G^-^) were separated using a BD FACSAria II cell sorter and collected into HBSS containing 1% BSA. Cells were checked for number and viability with 1:1 Trypan Blue under a brightfield microscope. Cells were mixed in the following ratio to reach a final total of 10,000: epithelial cells at 45%, endothelial cells at 45%, endothelial cells at 10%. This was performed separately for samples collected from each individual mouse. Following this, samples for each experimental group were mixed together to create a mixture of cells from two individual biological replicates with a final total of 20,000 cells. The cells were processed for single cell barcoding using the 10x Chromium platform and a library prepared for each sample using the 10X Genomics single-cell RNA-seq 3’ V3.1 kit. Sequencing was performed on the NovaSeq 2×150bp platform to a depth of 350 million reads per sample.

**Table 5.** Antibodies used for fluorescence activated cell sorting (FACS)

| Antibody | Clone | Supplier | Cat # | Dilution | RRID |
| --- | --- | --- | --- | --- | --- |
| Rat CD11b APC/Fire 750 | M1/70 | Biolegend | 101262 | 1:200 | AB_2572122 |
| Rat CD31 PE-Cy7 | MEC13.3 | Biolegend | 102523 | 1:800 | AB_2572181 |
| Rat CD326 PE | G8.8 | Biolegend | 118205 | 1:4000 | AB_1134176 |
| Rat CD45 BV510 | 30-F11 | Biolegend | 103138 | 1:200 | AB_2563061 |
| Rat F4/80 APC | BM8 | Invitrogen | 17-4801-82 | 1:200 | AB_2784648 |
| Rat Ly6G FITC | 1A8 | Biolegend | 127606 | 1:200 | AB_1236494 |
| Mouse SiglecF FITC | REA798 | Miltenyi | 130-112-178 | 1:200 | n/a |

### scRNA-seq pre-processing

Pre-processing of the data was performed using Seurat v4.0 according to the workflow suggested by the Satija lab [71]. First, ambient RNA was identified by comparing the raw and filtered matrices and contaminant RNA estimated using the SoupX package [72]. Adjusted matrices were then imputed into the R environment (v3.1) and analysed using Seurat. Normalisation was performed using regularised negative binomial regression via the SCTransform function, including the removal of confounding variation arising from mitochondrial mapping percentage [71]. Doublets were identified and removed using the “Doublet Finder” package via artificial next nearest neighbour analysis [73]. Batch effect correction was assessed using the Harmony package [74]. Identification of highly variable genes, next nearest neighbour, clustering functions and UMAP visualisation were all performed using the Seurat package. Marker genes per identified subpopulation were found using the findMarker function of the Seurat pipeline.

### SMG macrophage gene signature comparison

Single-cell RNA seq filtered matrix files from microglia (GSM3270885), colon (GSM3270887), spleen (GSM3270893), alveolar (GSM3270891) and Kupffer cell (GSM3270889) macrophages derived from Cre^-^ *Zeb2^fl/fl^ (*wildtype) mice were obtained from GEO Accession (GSE117079ID). Matrices were treated as above, with SCTransformation, doublet removal and batch effect correction before marker identification based on tissue of origin.

### Statistical tests

Normal distribution was assessed using the D’Agostino-Pearson omnibus test. Data were analysed for statistical significance using Student’s *t*-test (2 groups) or one-way ANOVA (multiple groups) with post-hoc testing performed using Tukey *Q* test (GraphPad Prism). For multiple testing we used a false discovery rate of 0.05. All graphs show the mean +standard deviation (SD), as indicated in the figure legends. Statistical tests used for each experiment are also indicated in all figure legends.

## Acknowledgements

The authors would like to thank Dr Andrea Corsinotti (CRM Multi-omics Facility), Dr Fiona Rossi and the CRM Flow Cytometry Facility, the CIR Flow Cytometry Facility, Dr Matthieu Vermeren and the CRM Imaging Facility, the CRM tissue culture facility, and the CRM and LFR animal facilities. JM is funded by UKRI/MRC grant MR/W004763/1 and the University of Edinburgh Chancellor’s Fellowship studentship; GRJ is funded by a Wellcome Trust Clinical Career Development Fellowship (220725/Z/20/Z); SE, EM and LH are funded by Wellcome Trust grant 108906/Z/15/Z; CR is funded by UKRI/MRC grant MR/S005544/1; NM is funded by UKRI/MRC grant MR/S000763/1 and UKRI/BBSRC grant BBS/E/D/10002071; ZL is funded by National Natural Science Foundation of China (31900630 and 32070880); MMP and RG are funded by The Kennedy Trust for Rheumatology Research and RG is funded also by a Chancellor’s Fellowship from the University of Edinburgh; CCB is funded by a Sir Henry Dale Fellowship jointly awarded by the Wellcome Trust and by the Royal Society (Grant number 206234/Z/17/Z); EE is funded by UKRI/MRC grant MR/S005544/1 and by a Chancellor’s Fellowship from the University of Edinburgh. CCB & EE have additional joint funding from the Wellcome Trust/ISSF3 Strategic Funds, The Royal Society (RGS\R2\202277) and the MRC Neuroimmunology Award Scheme (MR/W004763/1). The authors would like to thank Professor Stuart Forbes and Sue Pearson-Craven for critical reading of the manuscript. For the purpose of open access, the authors have applied a Creative Commons Attribution (CC BY) licence to any Author Accepted Manuscript version arising from this submission.

## Author Contributions

JM designed and performed experiments, analysed data and produced figures; GRJ analysed data, produced figures and co-wrote the manuscript; SE performed experiments and analysed data; EM, MMP, CR, LH and AJ performed experiments; ZL and FG originally generated and provided the *Ms4a3* mice, performed experiments and analysed data; CS provided the *Csf2ra^−/−^*mice; BB provided the *Il34^LacZ/LacZ^* mice; CP generated and provided the *Csf1*^FIRE/FIRE^ mice and provided edits to the manuscript; NM originally provided *Mafb^Cre^* and *Cx3cr1^LSL-DTR^*mice and provided edits to the manuscript; MB performed experiments, analysed data, produced figures and provided advice on experiments, interpretation and the manuscript; RG performed experiments, analysed data, produced figures, obtained funding, and provided advice on experiments, interpretation and the manuscript; CB and EE co-conceptualised the project, obtained funding, co-supervised the project, designed and performed experiments, analysed data, produced figures and co-wrote the manuscript.

## Supplementary Data

**Figure S1 – relates to Figure 1.**
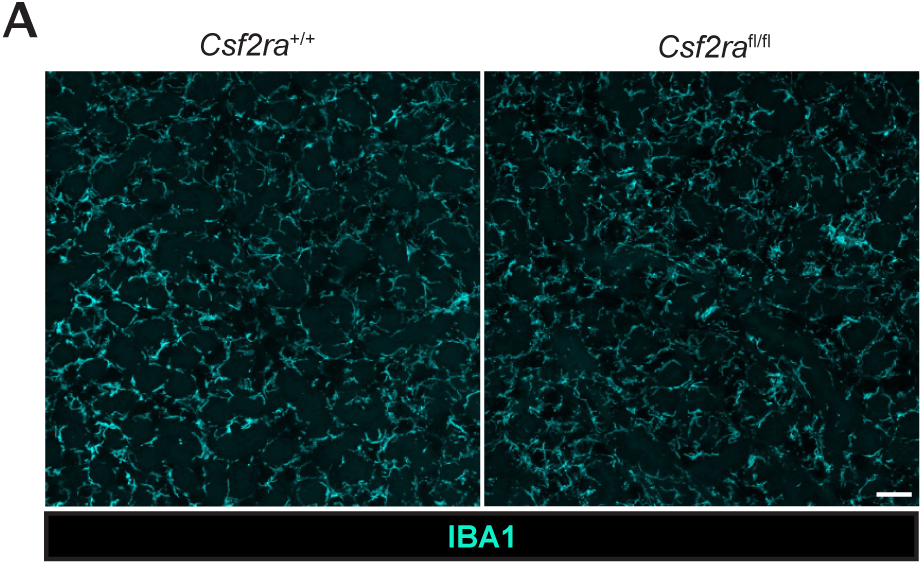
**A.** Representative expression of IBA1 in SMG tissue of *Csfr2a^+/+^* and *Csfr2a^fl/fl^* mice. Scale bar = 50μm.

**Figure S2 – relates to Figure 2.**
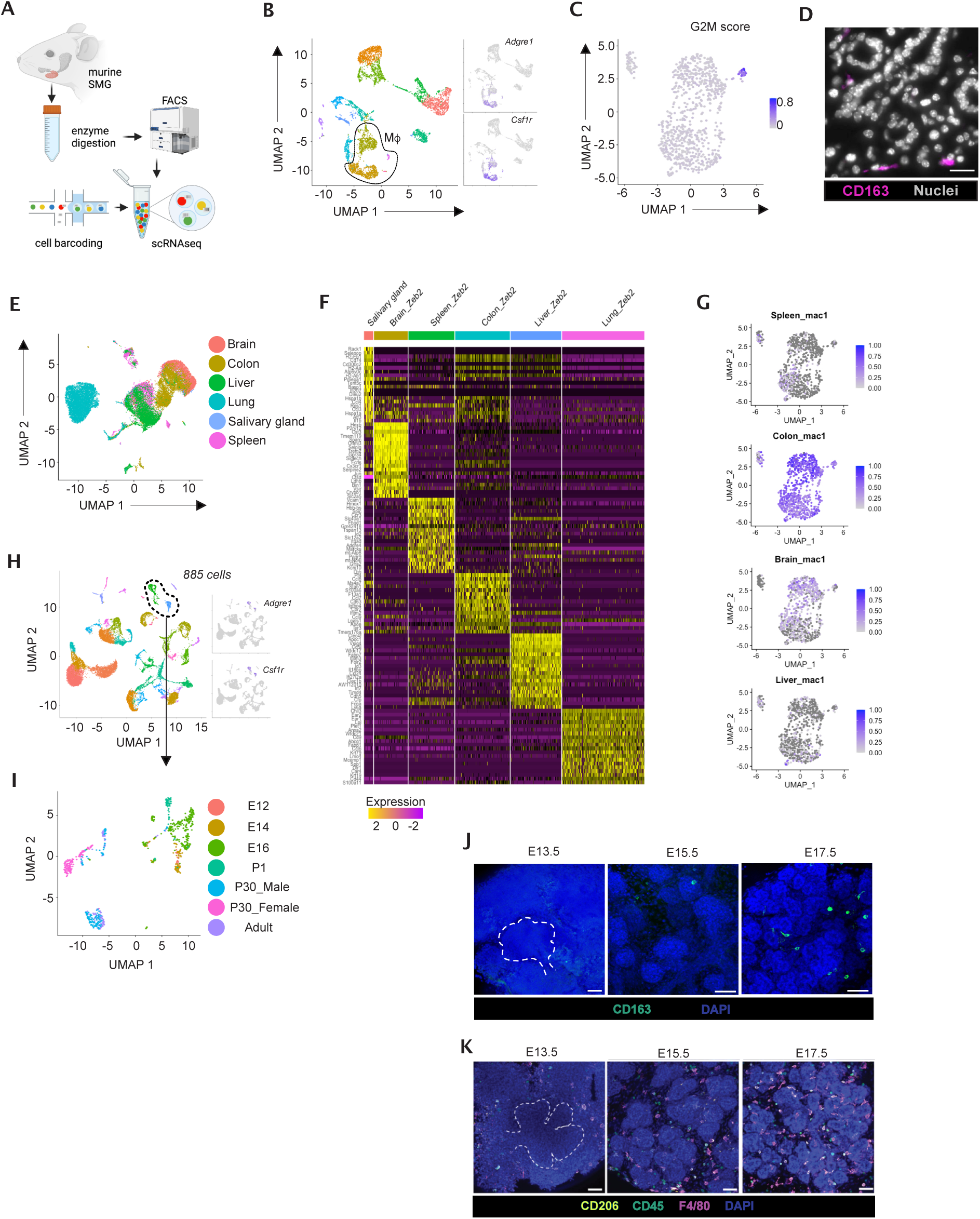
**A.** Schematic of scRNA-seq pipeline **B.** Feature plot showing expression of *Adgre1* and *Csf1r* to identify macrophages amongst epithelial and endothelial clusters. **C.** Feature plot showing G2/M phase, marker expression score across the macrophage population. **D.** Representative expression of CD163 in SMG tissue of unmanipulated adult C57BL/6 mice. Scale bar = 10μm. **E.** UMAP plot of comparative scRNA-seq gene expression analysis of SMG macrophages with brain microglia, colonic macrophages, lung alveolar macrophages, splenic red pulp macrophages and liver Kupffer cells from [28]. **F.** Heatmap showing the signature genes for SMG macrophages, brain microglia, colonic macrophages, lung alveolar macrophages, splenic red pulp macrophages and liver Kupffer cells **G.** Core macrophage gene expression profiles obtained from splenic, microglial and *Cd74^hi^*colon macrophages from single-cell data from [28] were used to generate a comparative quantification of single cell signature expression of the core profiles in SMG macrophages. **H.** UMAP plot of gene expression from existing scRNA-seq atlas of murine SMG development [30] containing E12, E14, E16, P1, P30 (male and female) and adult (10 months of age) datasets from C3H mice with feature plots of *Csf1r* and *Adgre1* expression **I.** UMAP plot of gene expression of *Csf1r / Adgre1* expressing cells from (H) split by SMG developmental stage [30]. **J.** Representative expression of CD163 in wholemount SMG tissue from unmanipulated embryonic (E13.5, E15.5, E17.5) C57BL/6J mice. Scale bars = 25mm. Dashed white line outlines SMG within the mesenchyme at E13.5. **K.** Representative expression of CD45, F4/80 and CD206 in wholemount SMG tissue from unmanipulated embryonic (E13.5, E15.5, E17.5) C57BL/6J mice. Scale bars = 25μm. Dashed white line outlines SMG within the mesenchyme at E13.5.

**Figure S3 – relates to Figure 4.**
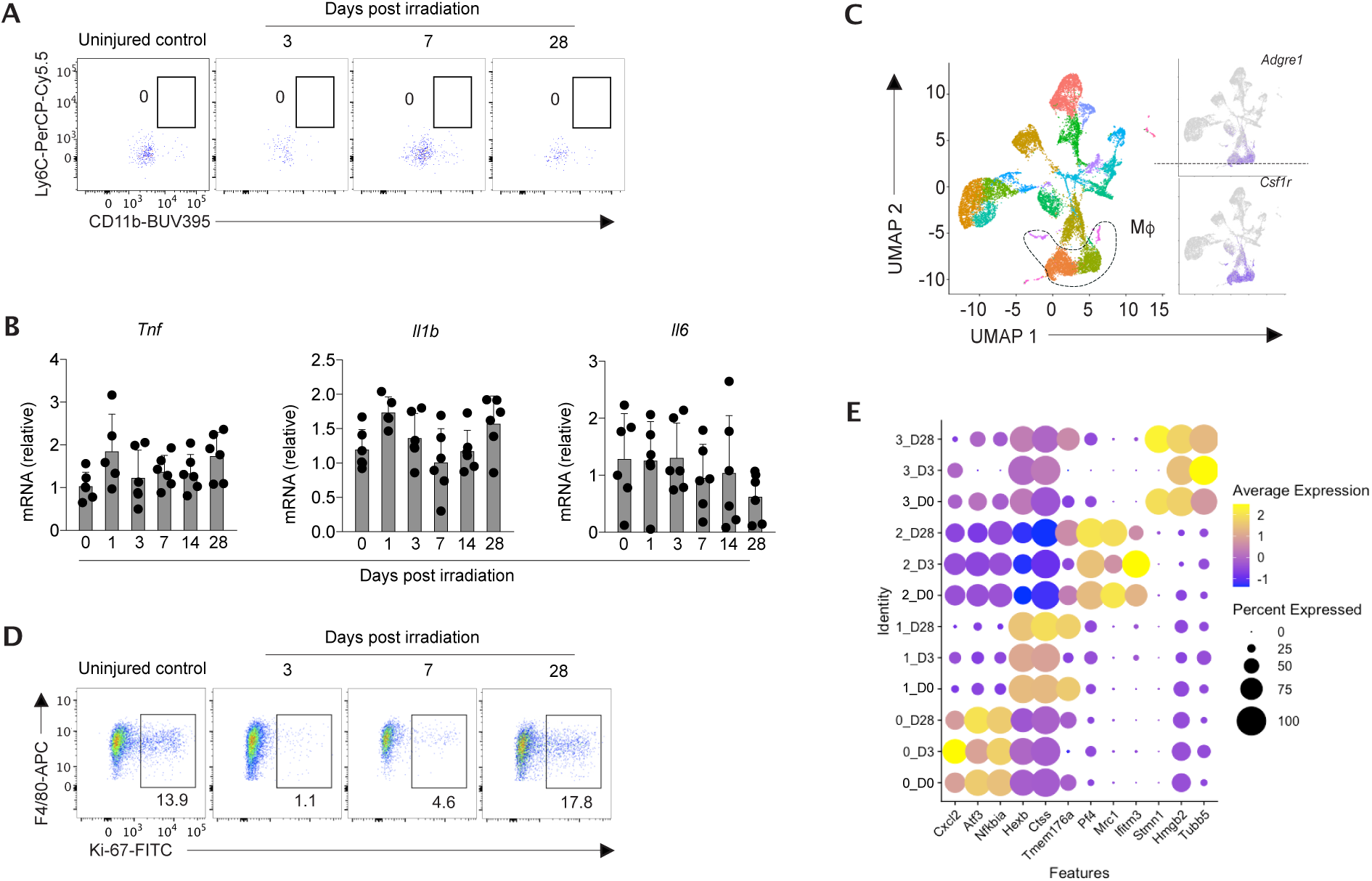
**A.** Representative expression of Ly6C and CD11b by F4/80^−^CD45^+^ cells from SMG at 3, 7 or 28 days post radiation or from unmanipulated mice. Data are from one of two independent experiments performed. **B.** qPCR analysis of *Tnf*, *Il1b* and *Il6* mRNA in total SMG tissue at the indicated time points following radiation induced injury. Data are normalised to mRNA levels in unmanipulated naïve (d0) SMG tissue. Data are from 3-6 mice pooled from 2 independent experiments. **C.** UMAP dimensionality reduction analysis of scRNA-Seq data from 14,855 cells containing macrophages, epithelia and endothelia from SMG of unmanipulated mice or mice irradiated 3 or 28 days earlier. Feature plots showing expression of *Adgre1* (encoding F4/80) and *Csf1r* to identify macrophages (Mf). **D.** Representative expression of Ki67 F4/80^+^CD11b^lo^ macrophages from SMG at 3, 7 or 28 days post radiation or from unmanipulated mice. Data are from one of two independent experiments performed. **E.** Bubble plot showing expression of selected genes by macrophage clusters from unmanipulated (D0) or at day 3 (D3) or day 28 (D28) following radiation treatment.

**Figure S4 – relates to Figure 5.**
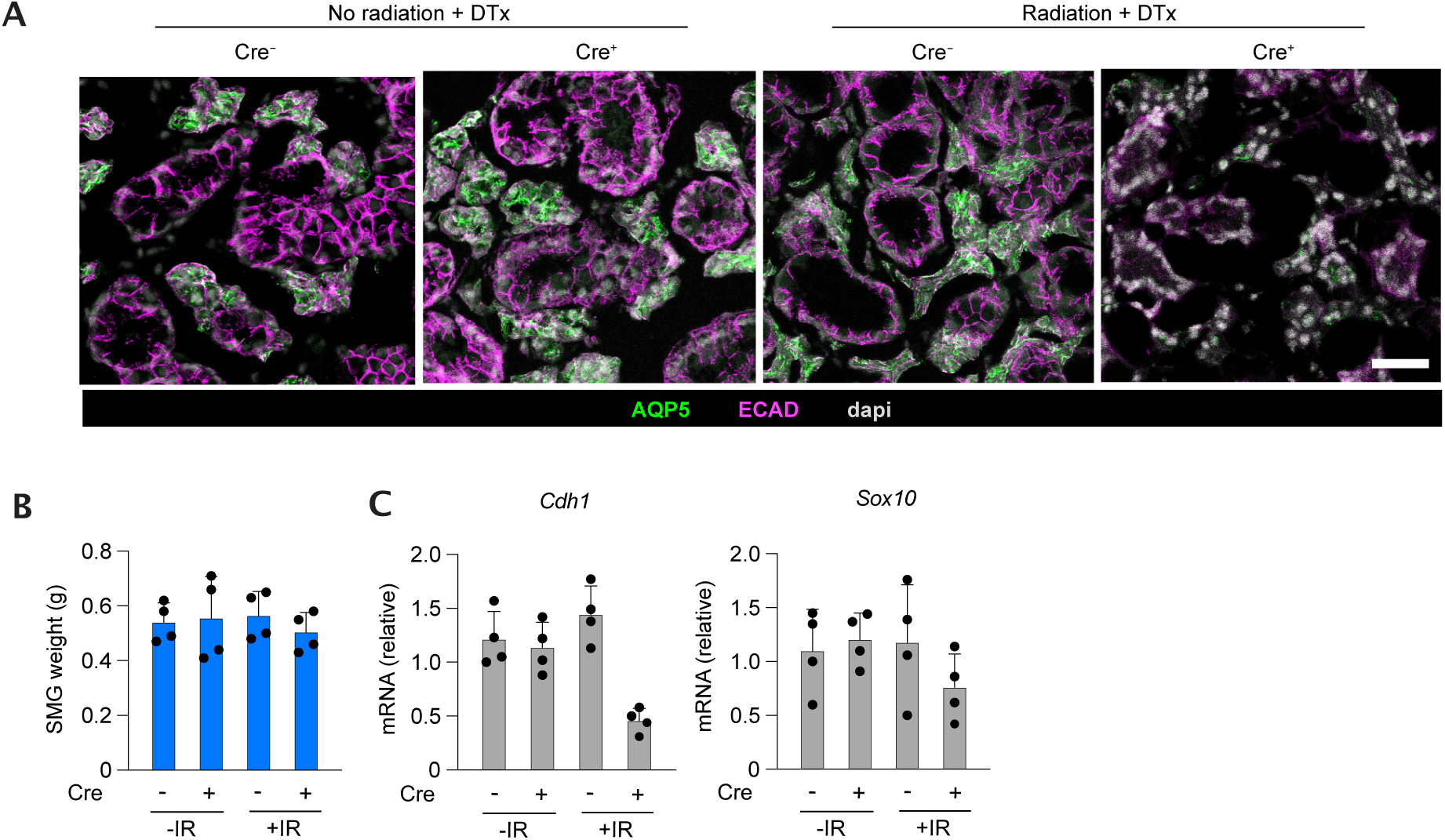
**A.** Additional immunofluorescent images of SMG stained for AQP5 and ECAD in *Mafb*^Cre/+^.*Cx3cr1*^LSL-DTR/+^ mice or *Mafb*^+/+^.*Cx3cr1*^LSL-DTR/+^ littermates exposed to 0Gy or 10Gy irradiation and administered diphtheria toxin (DTx) or saline from day 17 onwards, every 2 days and analysed 28 days after irradiation. Scale bar = 25μm. **B.** Quantification of single SMG weight of *Mafb*^Cre/+^.*Cx3cr1*^LSL-DTR/+^ mice or *Mafb*^+/+^.*Cx3cr1*^LSL-DTR/+^ littermates exposed to 0Gy or 10Gy irradiation and administered diphtheria toxin (DTx) or saline from day 17 onwards, every 2 days and analysed 28 days after irradiation. Data represent 1 SMG per mouse and are from 4 mice per group. **C.** qPCR analysis of *Cdh1 and Sox10* mRNA in total SMG tissue in *Mafb*^Cre/+^.*Cx3cr1*^LSL-DTR/+^ mice or *Mafb*^+/+^.*Cx3cr1*^LSL-DTR/+^ littermates exposed to 0Gy or 10Gy irradiation and administered diphtheria toxin (DTx) or saline from day 17 onwards, every 2 days and analysed 28 days after irradiation. Data are normalised to mRNA levels in SMG tissue *Mafb*^+/+^.*Cx3cr1*^LSL-DTR/+^ littermates exposed to 0Gy. Data are from 4 mice per group. *p<0.05, **p<0.01 (One-way ANOVA followed by Tukey *Q* post-hoc test).

